# Accurate inference of tree topologies from multiple sequence alignments using deep learning

**DOI:** 10.1101/559054

**Authors:** Anton Suvorov, Joshua Hochuli, Daniel R. Schrider

**Affiliations:** Department of Genetics, University of North Carolina at Chapel Hill, Chapel Hill, NC, USA; Biological & Biomedical Sciences Program, University of North Carolina at Chapel Hill, Chapel Hill, NC, USA

**Keywords:** supervised machine learning, convolutional neuronal network, phylogenetics

## Abstract

Reconstructing the phylogenetic relationships between species is one of the most formidable tasks in evolutionary biology. Multiple methods exist to reconstruct phylogenetic trees, each with their own strengths and weaknesses. Both simulation and empirical studies have identified several “zones” of parameter space where accuracy of some methods can plummet, even for four-taxon trees. Further, some methods can have undesirable statistical properties such as statistical inconsistency and/or the tendency to be positively misleading (i.e. assert strong support for the incorrect tree topology). Recently, deep learning techniques have made inroads on a number of both new and longstanding problems in biological research. Here we designed a deep convolutional neural network (CNN) to infer quartet topologies from multiple sequence alignments. This CNN can readily be trained to make inferences using both gapped and ungapped data. We show that our approach is highly accurate on simulated data, often outperforming traditional methods, and is remarkably robust to bias-inducing regions of parameter space such as the Felsenstein zone and the Farris zone. We also demonstrate that the confidence scores produced by our CNN can more accurately assess support for the chosen topology than bootstrap and posterior probability scores from traditional methods. While numerous practical challenges remain, these findings suggest that deep learning approaches such as ours have the potential to produce more accurate phylogenetic inferences.

## Introduction

Reconstructing the historical relationships among biological sequences remains one of the most important challenges in evolutionary biology. In order to reconstruct a tree it is first necessary to acquire orthologous (Koonin 2005) and/or paralogous (Hellmuth, et al. 2015) sequences from a sample of species or populations. Generally, the next step involves producing a multiple sequence alignment (MSA) in order to compare orthologous sequence characters, although in recent years a number of methods that bypass explicit orthology and alignment searches have been proposed (Hohl and Ragan 2007; Bonham-Carter, et al. 2013). Finally, any one of a plethora of tree reconstruction methods can be employed to infer a tree from an MSA. Determining the topology of a tree has been shown to be NP-hard for at least some formulations of the problem (Roch 2006), and the space of possible solutions grows double-factorially with increasing numbers of sequences (Felsenstein 1978b). We are thus limited to methods that utilize various heuristics and/or make assumptions about the process of sequence evolution in order to efficiently traverse tree space, including maximum parsimony (MP: Farris 1970; Fitch 1971), distance-based approaches such as neighbor joining (NJ: Saitou and Nei 1987), maximum-likelihood (ML: Felsenstein 1981) and Bayesian inference (BI: Rannala and Yang 1996; Li, et al. 2000; Huelsenbeck and Ronquist 2001). Although none of these methods are guaranteed to produce the true tree topology, one can evaluate and compare their effectiveness.

Supervised machine learning algorithms represent an alternative framework for drawing inferences from data, including sequence data. These methods have been successfully applied to various areas of biological research (Tarca, et al. 2007), and have recently been introduced to evolutionary biology—population genetics in particular (reviewed in Schrider and Kern 2018). Briefly, given some multivariate observation ***x*** that is associated with a response variable ***y***, supervised machine learning seeks to create a function *f*(***x***) that predicts ***y***. This is achieved through a process called training, wherein a “training set” of observations with known response variables are examined by an algorithm that creates and tunes the function *f* to minimize the disparity between *f*(***x***) and ***y*** on this set. Typically, ***x*** is summarized by a vector of values, or features. Convolutional neural networks (CNNs: Lecun, et al. 1998), a class of deep learning algorithms (Goodfellow, et al. 2016), are able to learn to extract informative features from data arranged in a matrix or higher-dimensional tensor in order to make a prediction; thus CNNs do not require a predefined feature vector. CNNs have proved extremely effective in analyzing image data (Krizhevsky, et al. 2012) and have recently been shown to achieve impressive accuracy on a number of tasks in population genetic inference when applied to MSAs (Chan, et al. 2018; Flagel, et al. 2018). CNNs and other deep learning methods may thus prove to make sizeable gains in a number of inference tasks involving biological sequence data.

Here we propose a novel approach applying CNNs to phylogenetic inference. We show that CNNs can be trained to extract phylogenetic signal from an MSA and use it to accurately reconstruct unrooted gene tree topologies (i.e. here our ***y*** is a tree topology) when given an alignment of four sequences. We adopt a classification approach, wherein the CNN is trained to discriminate among discrete data classes which in this case correspond to the three possible tree topologies. We find that our CNN approach is highly accurate on simulated data, can naturally include indel information, is fairly robust to classical biases affecting tree reconstruction and has several other benefits over existing methods. Below we describe our methodology and the CNN’s performance in detail before concluding with a brief discussion of limitations, potential extensions and outstanding practical challenges for adapting deep neural networks to phylogenomics.

## Methods

### MSA Simulations

For the 4-taxon case there exist exactly three possible unrooted tree topologies: ((A,B),(C,D)), ((A,C),(B,D)) and ((A,D),(B,C)) in Newick notation. We simulated an equal number of MSAs from each of these topologies using INDELible (Fletcher and Yang 2009), which allows for the inclusion of indels. The MSA length was set to 1000 sites unless stated otherwise. In order to construct a training set consisting of trees with a variety of branch length configurations we sampled uniformly from a “branch-length space” (BL-space) that encompasses a wide array of branch lengths and nodal rotations (see **Supplementary Text** and **Supplementary Fig. S1**). Tree branch lengths are expressed in expected number of substitutions per site.

With the tree topology and branch lengths in hand, sequences were generated under a randomly assigned Markov model of nucleotide substitution using INDELible. We used five nested models of increasing complexity: JC (Jukes and Cantor 1969), TIM, TIMef (Posada 2003), GTR (Tavaré 1986) and UNREST (Yang 1994) accommodating rate heterogeneity along the sequence by drawing the shape parameter of a continuous gamma distribution (+Γ) from *U*(0,4). We also allowed the proportion of invariable sites (*p*_inv_) in an MSA to vary, drawing this parameter (+I) from *U*(0,1). We note that the average expected substitutions per site (i.e. the branch length) is specified for variable sites only, not for the entire MSA. For example, when simulating an MSA with *p*_inv_ = 0.5, the specified branch lengths refer only to the 50% of sites that are free to vary while the remaining sites have branch lengths of zero. Substitution rate parameters and nucleotide frequencies were drawn from *U*(0,3) and a four-category Dirichlet distribution (**α** = 30), respectively. Finally, when including indels in the simulation we used identical fixed insertion and deletion rates (*λ*_I_ = *λ*_D_ = 0.01) and drew indel lengths from a Zipf distribution with *a* = 1.5, similar to empirical estimates (Vialle, et al. 2018), and a maximum indel size of 50. For each topology class we simulated either 5×10^4^, 1.5×10^5^ or 3×10^5^ MSAs for training and 1.5×10^4^ MSAs for the validation and test datasets. Additionally, we simulated test sets consisting of 3000 trees from each of thirteen regions of the tree-shape space that may potentially cause biased or more erroneous tree topology inference, including the Farris and Felsenstein zones that have long been known to negatively affect performance of classical methods (Swofford, et al. 2001). The simulation parameters for each of these were the same as those stated above with branch lengths drawn from the distributions shown in **Table 1**. Additionally we simulated trees within so called “twisted” Farris zone (McTavish, et al. 2015) with asymmetrical branch lengths as well as trees from different regions that may potentially cause tree estimation to be less accurate. Throughout this paper we denote sister branches or their branch lengths on the left side of the tree as B_1_ and B_2_, those on the right side as B_3_ and B_4_, and the internal branch as B_5_. The overall pipeline is summarized in **Figure 1**.

**Figure 1:**
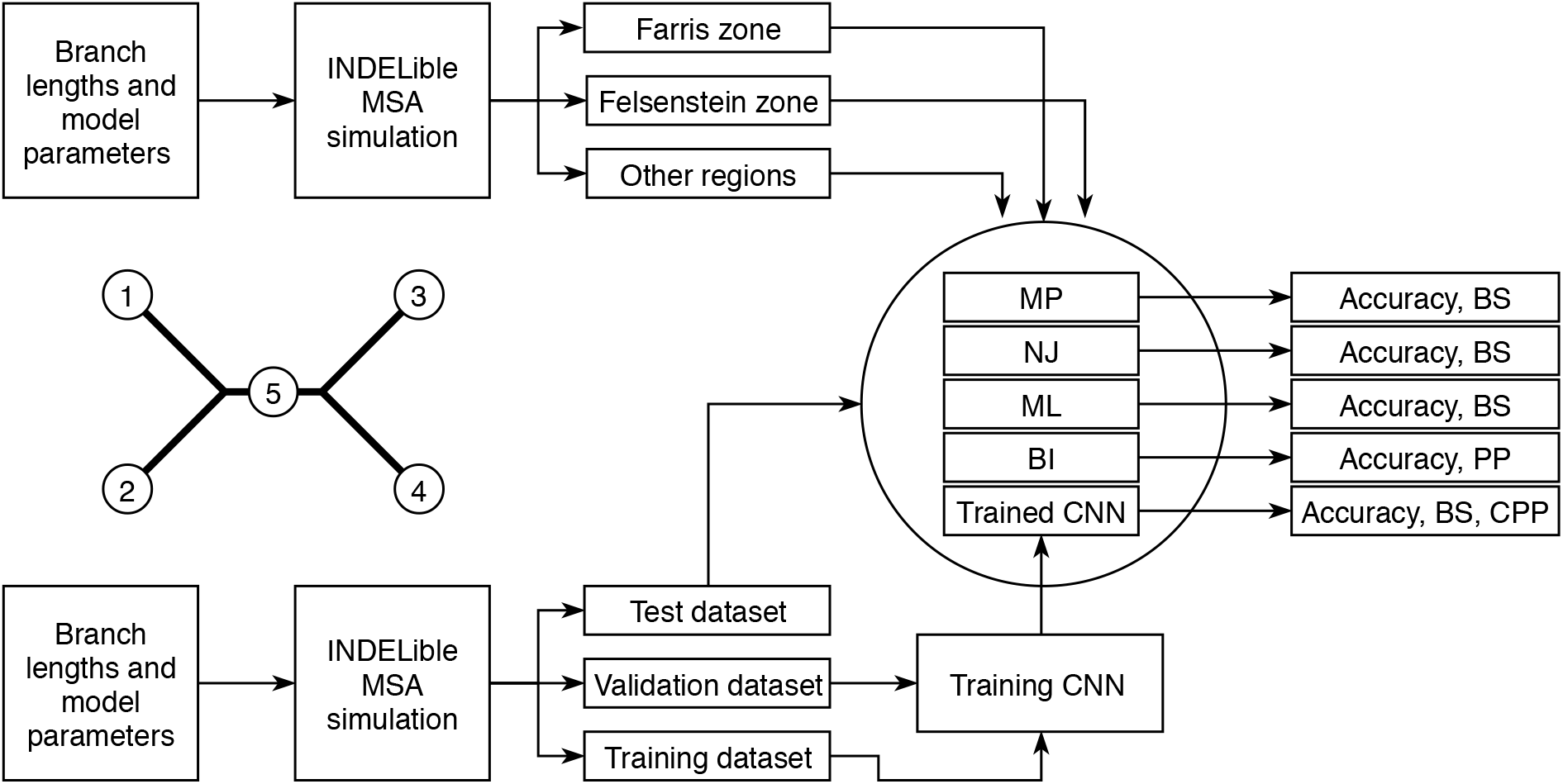
Summary of the simulation and testing procedures. The top branch of the diagram represents the steps that were used to simulate datasets with various branch length configurations (**Table 1**) to test each method’s performance. The bottom branch of the diagram represents the steps that were used to simulate training and validation datasets to train the CNN, and the initial test dataset was used to estimate accuracy for each method. The training, validation and test datasets were simulated using the procedure described in **Supplementary Text**. MSA=multiple sequence alignment, MP=maximum parsimony, NJ=neighbor-joining, ML=maximum likelihood, BI=Bayesian inference, CNN=convolutional neuronal network, BS=bootstrap, PP=posterior probabilities, CPP=class membership posterior probability. The tree shows the branch labelling convention that was used throughout the paper, where numbers in circles correspond to branches B_1_, B_2_, B_3_, B_4_ and B_5_.

**Table 1:** Branch-length distributions for test sets designed to assess accuracy and the presence of bias in topology inference. B_1_ and B_2_ and B_3_ and B_4_ denote sister branches on the left and right sides of a tree respectively whereas B_5_ denotes the internal branch. *TExp* denotes truncated exponential distribution. *Beta* denotes beta distribution. A mixture of beta components with equal weights (i.e. w_1_ = w_2_ = w_3_ = 1/3) was used to generate the space from which samples were drawn as described in the **Supplementary Text**. *U* denotes uniform distribution. B_1_+B_5_ is a sum of two uniform random variables which follows the Irwin-Hall distribution.

| Region | $B_1$ | $B_2$ | $B_3$ | $B_4$ | $B_5$ |
| --- | --- | --- | --- | --- | --- |
| BL-space | Sampled from $w_1\text{Beta}(\alpha=\beta=0.1)+w_2\text{Beta}(\alpha=\beta=0.5)+w_3\text{Beta}(\alpha=\beta=1)$ as described in **Supplementary Text** | | | | |
| Truncated exponential | All *TExp*(10,0,0.5) |  |  |  |  |
| Farris zone | $U(0.1,0.5)$ | $U(0.1,0.5)$ | $U(0,0.05)$ | $U(0,0.05)$ | $U(0,0.05)$ |
| Twisted Farris zone | $U(0.1,0.5)$ | $B_1+B_5$ | $B_5$ | $2B_5$ | $U(0,0.05)$ |
| “Extended” Farris zone | $U(0.1,0.5)$ | $U(0.1,0.5)$ | $U(0,0.05)$ | $U(0,0.05)$ | $U(0,0.5)$ |
| Felsenstein zone | $U(0.1,0.5)$ | $U(0,0.05)$ | $U(0.1,0.5)$ | $U(0,0.05)$ | $U(0,0.05)$ |
| “Extended” Felsenstein zone | $U(0.1,0.5)$ | $U(0,0.05)$ | $U(0.1,0.5)$ | $U(0,0.05)$ | $U(0,0.5)$ |
| Long branches | $U(0.1,0.5)$ | $U(0.1,0.5)$ | $U(0.1,0.5)$ | $U(0.1,0.5)$ | $U(0,0.5)$ |
| Extra-long branches | $U(0.5,1)$ | $U(0.5,1)$ | $U(0.5,1)$ | $U(0.5,1)$ | $U(0,1)$ |
| Single long branch | $U(0.1,0.5)$ | $U(0,0.05)$ | $U(0,0.05)$ | $U(0,0.05)$ | $U(0,0.5)$ |
| Short branches | $U(0,0.05)$ | $U(0,0.05)$ | $U(0,0.05)$ | $U(0,0.05)$ | $U(0,0.5)$ |
| Extra-short branches | All $U(0,0.01)$ | | | | |
| Single short branch | $U(0,0.05)$ | $U(0.1,0.5)$ | $U(0.1,0.5)$ | $U(0.1,0.5)$ | $U(0,0.5)$ |
| Short internal branch | $U(0,0.5)$ | $U(0,0.5)$ | $U(0,0.5)$ | $U(0,0.5)$ | $U(0,0.05)$ |

The internal branch length (B_5_) for the Farris and Felsenstein zones was drawn from *U*(0,0.05) (**Table 1**). In order to investigate effect of internal branch length on each method’s performance in more detail, we also created variants of the Farris and Felsenstein zones, called “extended” where B_5_ was drawn from the interval *U*(0,0.5). Previous simulation studies suggest that methodological biases primarily arise when an internal branch appears to be very short, i.e. internal branch length ≤0.05 (Swofford, et al. 2001). However, since these studies followed overly simplified simulation approaches (e.g. MSA simulation under JC model only; Siddall 1998), this conclusion may not hold for MSA datasets generated using more complex simulation strategies that incorporate various substitution models, rate heterogeneity, varying proportions of invariant sites and indels as we do in this study.

### CNN Architecture and Training

This section describes the structure of our neural network and training procedure, and thus many of the concepts herein may be foreign to some readers. We refer those interested in an accessible overview of CNNs in the context of sequence alignments to Flagel et al. (2018). We implemented our CNN in Python using version 2.2.4 of the Keras API (https://keras.io/) with TensorFlow 1.12.0 (Abadi, et al. 2016) as the backend. To extract phylogenetically informative features from an MSA we used a 1D convolution-pooling strategy. We represented each MSA as a matrix with rows corresponding to sequences and columns to sites in the alignment, and with each character in the alignment encoded as a number (‘A’:0, ‘T’:1, ‘C’:2, ‘G’:3, ‘-’:4), and encoded the MSA’s associated topology as a numeric class label. Since indels result in MSAs of varying lengths and our CNN requires all input matrices to have equal size, shorter alignments were extended by “padding” the MSA with additional columns wherein each value was arbitrarily set to −15 until the MSA’s length matched that of the longest alignment in our simulated data. Our CNN consisted of eight convolution layers with ReLU activation (Nair and E. Hinton 2010) and each followed by an average-pooling step, one 1024-node fully connected layer with ReLU activation and an output layer with three nodes corresponding to the three possible topology classes. We used softmax as the activation function for our output layer in order to assign a class membership probability to each of the three classes. We used categorical cross-entropy as our loss (i.e. error) function. During training we used the adaptive moment estimation (Adam) optimizer (Kingma and Ba 2014).

We set the filter size for the first convolution operation to 4×1, reasoning that it would potentially capture a phylogenetic signal by striding across the MSA and examining one column of the alignment at a time. The total number of filters was set to 1024 for the first and second convolutional layer and set to 128 for the remaining convolutional layers. We decided upon 1024 filters because this number is greater than the number of possible columns in an MSA with four DNA sequences allowing gap characters, and we had decided (arbitrarily) to limit these values to powers of two. Additional network hyperparameters were chosen somewhat arbitrarily after assessing performance of several different network architectures. We also used average-pooling rather than the more common max-pooling operation given that Flagel et al. found this approach to be effective when applied to population genetic alignments (Flagel, et al. 2018). The first average-pooling operation size was set to 1×1 (i.e. no pooling), then all the subsequent convolution steps had equal sizes of 1×2 and pooling steps had equal sizes of 1×4, 1×4, 1×4, 1×2, 1×2 and 1×1. We had also experimented with fewer layers but noticed that “deeper” CNN architectures, i.e. with more convolutional layers had better accuracy. In order to avoid possible internal covariate shift and model overfitting we used batch normalization (Ioffe and Szegedy 2015) followed by dropout regularization with a rate of 0.2 for each convolution layer as well as for the hidden layer. During training in most of the cases we used a minibatch size of 150 MSAs. The detailed network architectures along with the training time are available in **Supplementary Text**. The training procedure was run for a number of iterations determined by the stopping criterion (improvement in validation loss of the current iteration over the best prior iteration < 0.001) and patience value (the number of consecutive iterations that must satisfy the stopping criterion before halting training). We observed that the results of our training runs were fairly stable, with a given CNN architecture consistently achieving the same final accuracy on the validation set. To further examine performance during training, we visualized the surface of the loss function and training trajectories across the CNN’s parameter space (i.e. the possible values of all weights in the network) as described below. This revealed that there appear to be many readily reachable local minima of the loss function that all have roughly equivalent values (**Supplementary Fig. S2**), lending further support to the notion that training results should be fairly stable, at least for the datasets and architectures examined here.

### Testing Procedures

We sought to compare the performance of CNN vs. conventional tree inference methods, namely MP, NJ, ML and BI using our simulated test datasets. MP reconstructions were performed using PHYLIP’s dnapars program v3.696 (Felsenstein 1989) in the interface provided by SeaView v4.7 (Gouy, et al. 2010). ML analysis was implemented in IQ-TREE v1.6.5 (Nguyen, et al. 2015) restricting model selection to only the models that were used to simulate MSAs, i.e. JC, TIM, TIMef, GTR and UNREST (equivalent to the 12.12 model in IQ-TREE) with +I and/or +Γ, where the continuous rate gamma distribution was approximated by discrete gamma with 8 categories (G8). For each MSA, ML search was performed exhaustively evaluation each of the three possible unrooted topologies. The NJ trees were generated using the distance function of the best-fit model in IQ-TREE. Note that ML and NJ approaches infer a rooted tree under the UNREST model, thus every such estimated tree was unrooted for further comparisons. Bayesian tree reconstruction was carried out in ExaBayes v1.4.1 (Aberer, et al. 2014) under default parameters starting with a random initial topology. Three independent MCMC chains were run for 10^6^ generations each with two coupled chains. Tree support was assessed by standard nonparametric bootstrap (Felsenstein 1985) for MP, NJ, ML and CNN, and by posterior split probability for BI. We also assessed tree support from the CNN by simply using the estimated posterior class membership probabilities emitted by the output layer. Then, for each method we calculated topological accuracy as the number of correctly returned topologies divided by the total number of inferences performed. See **Supplementary Text** for exact commands that were used to infer topologies.

## Results and Discussion

### CNNs are more Accurate than Standard Methods on Gapped and Ungapped MSAs within BL-space

Initially, we trained our CNN on gapped and ungapped training sets of total 15×10^4^ MSAs (each with 1000 sites, 5×10^4^ MSAs for each of our three possible topologies) and in each case for our initial evaluation we assessed our accuracy on an independent test set generated in the same manner as our training data. We compared the accuracy of our CNN on simulated test data (1.5×10^4^ MSAs per topology) to that of several other standard tree estimation methods, namely maximum parsimony (MP), neighbor joining (NJ), maximum likelihood (ML), and Bayesian inference (BI). Strikingly, the CNN was substantially more accurate than the other methods (McNemar test, *P* < 1×10^−10^ for each comparison with the CNN) when applied to gapped MSAs (**Fig. 2**). However, this was not the case when testing methods on ungapped MSAs: here accuracy of the CNN (0.765), which was in this case also trained on ungapped MSAs, was nearly identical to the best performing method, BI (accuracy = 0.765, McNemar test, *P* = 0.918). Nevertheless, the CNN’s accuracy improved with increasing amounts of training data and obtained higher accuracy (0.777; **Fig. 2**) than all other methods when trained on either 1.5×10^5^ or 3×10^5^ ungapped MSAs per topology. The difference with the next best method (BI) was not significant (McNemar test, *P* = 0.08) with 1.5×10^5^ training examples but significant (McNemar test, *P* = 0.013) with 3×10^5^ training examples. Although the results of this initial evaluation within our BL-space are encouraging, we must bear in mind that accuracy can vary within this space. Moreover, our BL-space was constructed in part to expose CNN to many branch-length configurations including some extreme configurations which one may not expect to commonly encounter in real data sets. Thus in the next section we test each method on a number of more narrowly defined tree branch-length configurations in order to gain a more detailed picture of our method’s strengths and weaknesses.

**Figure 2:**
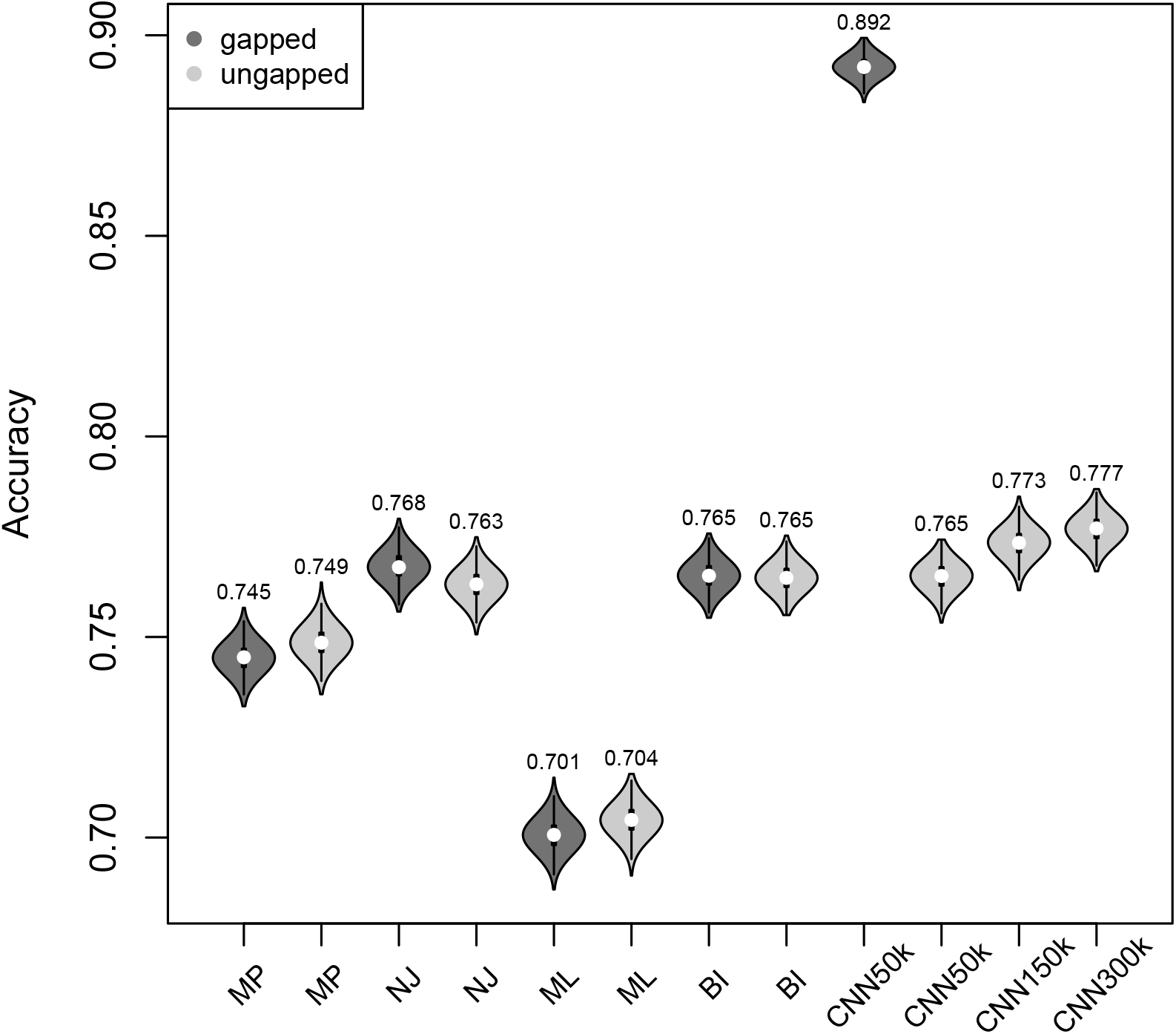
Accuracy of tree topology inference within the test set. The violin plots show the accuracy for different tree estimation methods achieved on gapped and ungapped (15000 MSAs each) datasets. The distribution shown in each violin plot is obtained by bootstrap resampling from the test dataset and calculating accuracy on each bootstrap replicate. The white dots represent the accuracy point estimate (across the complete test set) whose value is indicated above each violin. The number after CNN (i.e. 50k, 150k or 300k) indicates number of MSAs per topology that were used to in training. MP=maximum parsimony, NJ=neighbor-joining, ML=maximum likelihood, BI=Bayesian inference, CNN=convolutional neuronal network.

We also investigated the relationship between several simulation parameters and inference accuracy. When sequences are very closely related, the MSA may consist largely of invariant sites, which are phylogenetically uninformative. One of the parameters of our simulations was the fraction of sites that were not allowed to vary (*p*_inv_; see Methods). We found that for each method, misclassified MSAs from the gapped test set had a larger value of *p*_inv_ on average (Wilcoxon rank sum test, *P* < 1×10^−10^ for each standard method and *P* = 6.135×10^−5^ for the CNN), but this difference is far less pronounced for the CNN (**Fig. 3a**). However, in the case of the ungapped test set, MSAs misclassified by all methods including the CNN showed a similar bias toward larger values *p*_inv_ on average (Wilcoxon rank sum test, *P* < 1×10^−10^ for each method; **Fig. 3b**). The gamma shape parameter, *α*, which governs mutation rate heterogeneity across the MSA does not impact the CNN’s ability to infer the correct topology (Wilcoxon rank sum test, *P* = 0.91) whereas for all other methods misclassified MSAs had elevated values of *α* (Wilcoxon rank sum test, *P* < 1×10^−5^ for all standard methods; **Fig. 3c**). Again, in the case of ungapped data each method showed a similar bias, with larger *α* in misclassified versus correctly classified examples (Wilcoxon rank sum test, *P* < 1×10^−5^ for all methods; **Fig. 3d**). Together these results suggest that CNNs are more robust to higher values of *p*_inv_ and *α* than traditional methods for gapped MSAs. Additionally, we note that the DNA substitution model does not affect any method’s performance (Fisher exact test, *P* > 0.05 for all methods).

**Figure 3:**
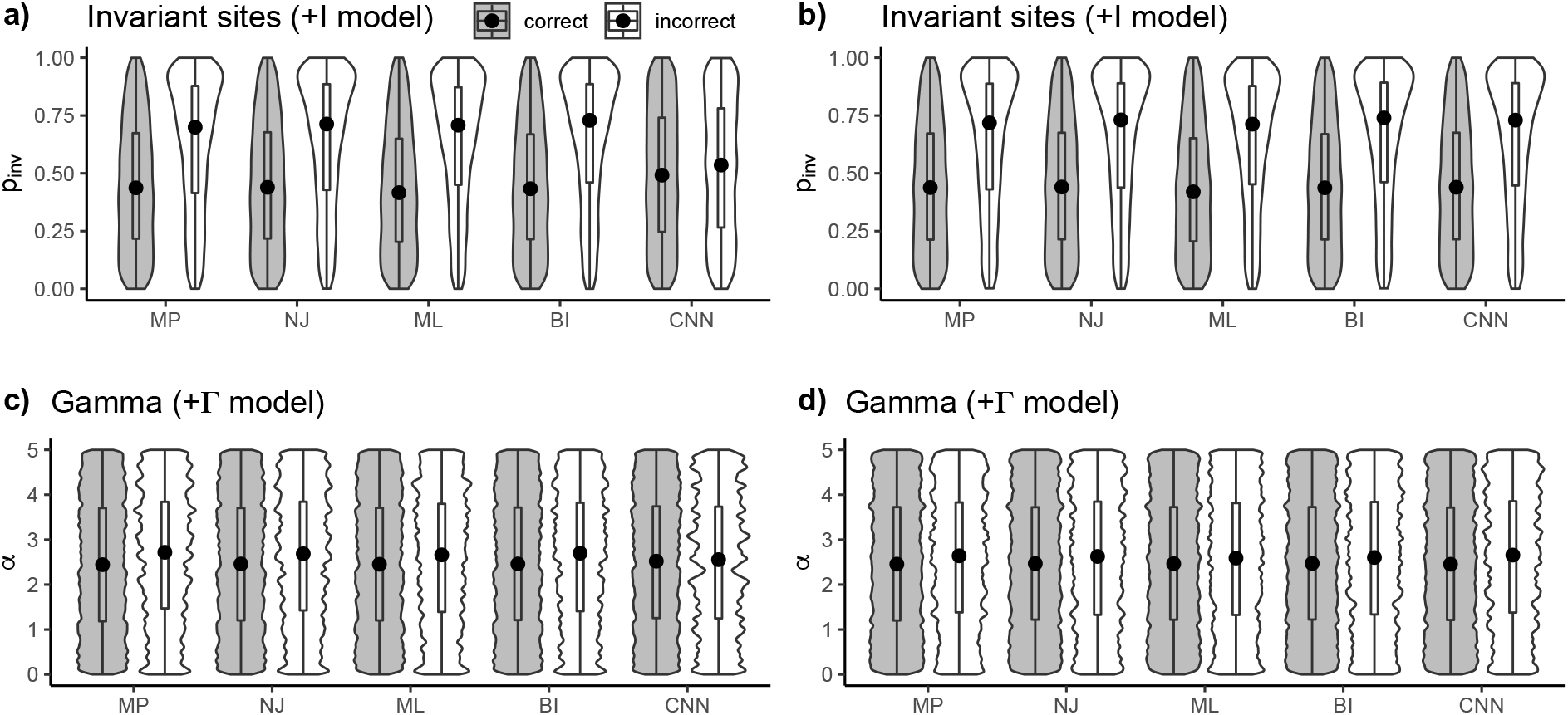
Differences in parameter values between correct and incorrect classifications for each method. The distribution of the parameter governing the fraction of invariable sites (*p*_inv_) for correctly and incorrectly inferred trees for a) gapped and b) ungapped MSAs. The distribution of gamma rate parameter *(α)* for correctly and incorrectly inferred trees for c) gapped and d) ungapped MSAs. The black dots of each violin plot indicate median of the distribution. These results were obtained using 15000 simulated MSAs from BL-space. MP=maximum parsimony, NJ=neighbor-joining, ML=maximum likelihood, BI=Bayesian inference, CNN=convolutional neuronal network.

Overall these results suggest that our CNN can effectively extract phylogenetic signal. This is especially so for the CNN trained on gapped MSAs, which easily outcompeted the other methods, all of which treat gaps as missing data. Despite their potential to provide unambiguous phylogenetic information (Rokas and Holland 2000; Dessimoz and Gil 2010) due to reduced homoplasy relative to substitutions (Ashkenazy, et al. 2014), approaches incorporating indels are controversial and not generally utilized in practice (Ogden and Rosenberg 2007; Warnow 2012). When our CNN is trained on gapped data it learns to detect phylogenetic signal from both indels and substitutions present in an MSA. To further examine this property of the gapped MSA CNN, we simulated MSAs of length 1000 using trees with branch lengths drawn from U(0,0.5) with no substitutions (i.e. *p*_inv_ = 1) allowing only indels. Under these conditions all standard methods fail to achieve consistency (Warnow 2012; Truszkowski and Goldman 2016), and MP, NJ, ML and BI all had accuracy of ~0.33 or lower—no better than random guessing. By contrast, the CNN’s accuracy on this set was 0.927. To test a scenario towards the opposite extreme wherein every site is informative, we simulated MSAs of length 1000 allowing all sites to vary (*p*_inv_ = 0) under the JC model, including indels, and using trees with all branch lengths drawn from U(0,0.5). Under these conditions all methods exhibited very high accuracy (MP: 0.962, NJ: 0.976, ML: 0.975, BI: 0.978 and CNN: 0.959).

These results further suggest that our CNN, when trained on gapped MSAs, is in principle able to incorporate indel information into its phylogenetic inference. Because the CNN implicitly constructs substitution and indel process models by examining the simulated training data, alignment error could impact inference in practice. This could be handled in one of two ways: 1) by incorporating alignment errors into the training set (e.g. by running a sequence aligner on training data prior to training), or 2) judging the reliability of indels within the MSA and masking particular sites or omitting entire MSAs appearing to be unreliable (e.g. following Castresana 2000; Misof and Misof 2009; Ashkenazy, et al. 2014; Fujimoto, et al. 2016). Another potential problem is model misspecification: accuracy could suffer if indel rates/lengths differ dramatically between simulated training data and real datasets. Future work will have to investigate strategies to train a network that is robust to such misspecification, perhaps by incorporating a variety of indel models into training; additional steps to train neural networks to be insensitive to the proclivities of empirical data may also be necessary (discussed in Concluding Remarks below).

### Topology Estimation Under Branch Length-Heterogeneity

There exist several regions in the branch-length parameter space where common methods fail to produce the correct topology. The bias known as long branch attraction (LBA) is the inability of MP (Felsenstein 1978a) and possibly Bayesian (Kolaczkowski and Thornton 2009; Susko 2015) methods to estimate the correct topology in presence of long nonadjacent branches (i.e. the Felsenstein zone; Huelsenbeck and Hillis 1993); these methods tend to erroneously place such long branches together and thus favor an incorrect topology (**Table 1; Figs. 4c and 5c**). Another region in the branch-length space, called the Farris zone (Siddall 1998), causes ML methods to exhibit reduced accuracy on topologies that have adjacent long branches and causes MP to correctly infer trees more often as a result of LBA which in this case biases it toward the correct topology (**Table 1; Figs. 4a and 5a**); we note that this bias may diminish when using longer alignments due to ML’s property of consistency (Swofford, et al. 2001). Additionally, we assessed the performance of all methods in the asymmetrical (“twisted”) Farris zone (**Table 1; Figs. 4b and 5b**), where a single incorrect topology is favored by NJ if the corrected distance function is convex (McTavish, et al. 2015). We also examined trees with a short internal branch (≤ 0.05) but external branches lengths varying along the interval [0,0.5] (**Table 1; Figs. 4d and 5d**).

**Figure 4:**
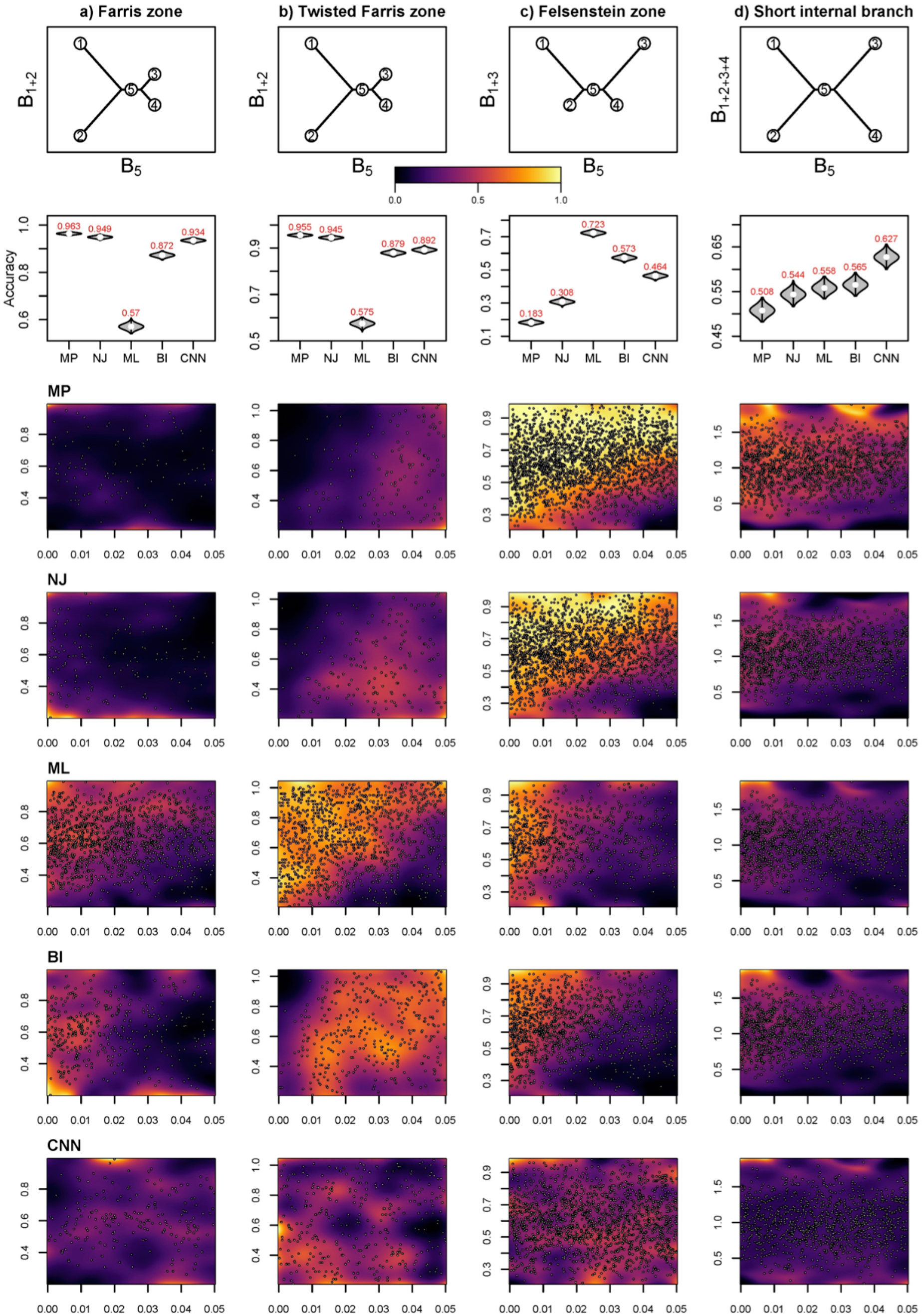
Accuracy within regions with elevated branch-length heterogeneity for gapped MSAs. The four columns summarize results in a) the Farris zone, b) the twisted Farris zone, c) the Felsenstein zone and d) trees with short internal branches. The top row illustrates these tree branch length configurations, and the second row shows accuracy of each method’s performance in each zone (with violins representing accuracy on bootstrap replicates from the test set). Panels three through seven show the locations (gray dots) and densities (heat map) of incorrectly inferred topologies for each method plotted in two-dimensional space with B5 length on the *x*-axis and B_1_+B_2_ (labeled as “B_1+2_”), B_1_+B_2_ (“B_1+2_”), B_1_+B_3_ (“B_1+3_”), B_1_+B_2_+B_3_+B_4_ (“B_1+2+3+4_”) on the *y*-axis for Farris, twisted Farris, Felsenstein and short internal branch regions respectively. See Table 1 for branch length simulation details. The color scale shows the estimated density of the incorrectly inferred topologies at each point in the branch-length parameter space, with brighter colors representing a greater density of errors. MP=maximum parsimony, NJ=neighbor-joining, ML=maximum likelihood, BI=Bayesian inference, CNN=convolutional neuronal network.

**Figure 5:**
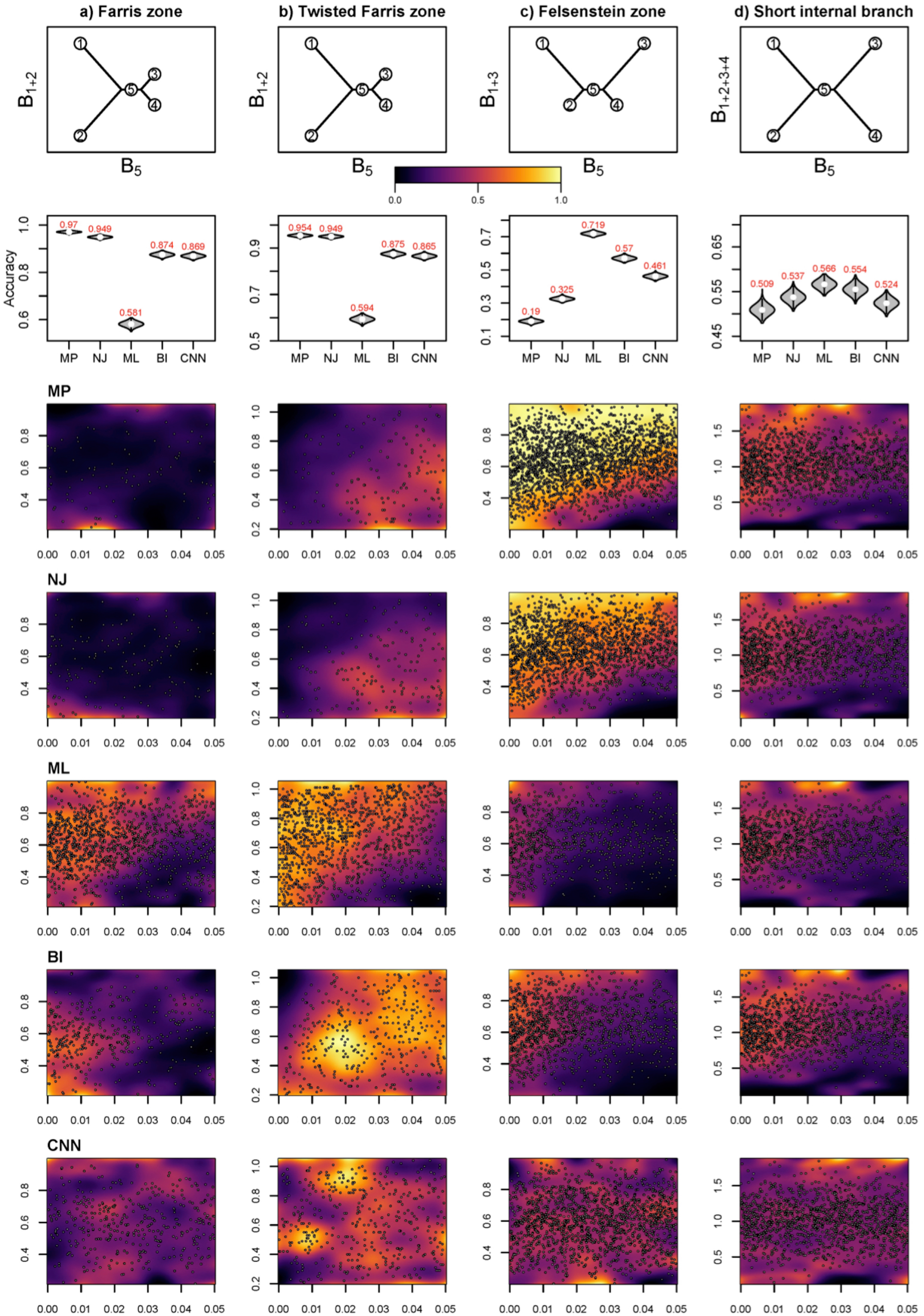
Accuracy within regions with elevated branch-length heterogeneity for ungapped MSAs. The four columns summarize results in a) the Farris zone, b) the twisted Farris zone, c) the Felsenstein zone and d) trees with short internal branches. The top row illustrates these tree branch length configurations, and the second row shows accuracy of each method’s performance in each zone (with violins representing accuracy on bootstrap replicates from the test set). Panels three through seven show the locations (gray dots) and densities (heat map) of incorrectly inferred topologies for each method plotted in two-dimensional space with B5 length on the *x*-axis and B_1_+B_2_ (labeled as “B_1+2_”), B_1_+B_2_ (“B_1+2_”), B_1_+B_3_ (“B_1+3_”), B_1_+B_2_+B_3_+B_4_ (“B_1+2+3+4_”) on the *y*-axis for Farris, twisted Farris, Felsenstein and short internal branch regions respectively. See Table 1 for branch length simulation details. The color scale shows the estimated density of the incorrectly inferred topologies at each point in the branch-length parameter space, with brighter colors representing a greater density of errors. MP=maximum parsimony, NJ=neighbor-joining, ML=maximum likelihood, BI=Bayesian inference, CNN=convolutional neuronal network.

We primarily focus on these four regions of known bias (**Figs. 4 and 5**) but also examined several other regions of potential bias on test sets of gapped (**Table 1; Supplementary Fig. S3**) and ungapped (**Table 1; Supplementary Fig. S4**) MSAs. The accuracy estimates for each region are summarized in **Table 2**. In most of these regions, our CNN exhibited accuracy roughly matching or exceeding that of the best performing traditional method, whereas each other method failed catastrophically in at least one of these regions (**Figs. 4 and 5, Supplementary Figs. S3, S4**). The two exceptions where the CNN exhibited markedly reduced accuracy were the Felsenstein zone (**Figs. 4 and 5; Table 2**) and the “extra-short branches” region where all branch lengths range vary uniformly along [0, 0.01] (**Table 2; Supplementary Figs. S3 and S4**). Specifically, in the Felsenstein zone the CNN’s accuracy is 0.464 (gapped) and 0.461 (ungapped), above the random expectation of 1/3, and well below that of ML (0.723 (gapped) and 0.719 (ungapped); **Figs. 4c and 5c**). However, CNN’s accuracy could be notably improved by adding additional MSAs simulated from the Felsenstein zone or the “extra-short branches” region: performing additional training iterations on 1000 MSAs per topology from these regions boosted the CNN’s accuracy to 0.753 for gapped MSAs and to 0.771 for ungapped MSAs for the Felsenstein zone, and to 0.882 (gapped) and 0.904 (ungapped) for the “extra-short branches” region respectively (**Table 2**). This suggests that it is possible to fix a known “blind spot” of the CNN by simply simulating additional relevant training data and performing more iterations of the training algorithm. The CNN shows superior accuracy in each of the remaining regions of potential bias (**Supplementary Fig. S3**) for gapped MSAs compared to all other methods. Overall, on ungapped data, the CNN’s performance roughly matched that of BI across zones (**Supplementary Fig. S4**), with the exception of the regions mentioned above. These two methods both exhibit acceptable but not exceptional accuracy in each of these different zones (**Table 2**), perhaps with the exception of the Felsenstein zone where all methods other than ML have low accuracy (≤ 0.57 versus 0.719 for ML), unless the CNN is provided with additional training data as described above.

**Table 2:**
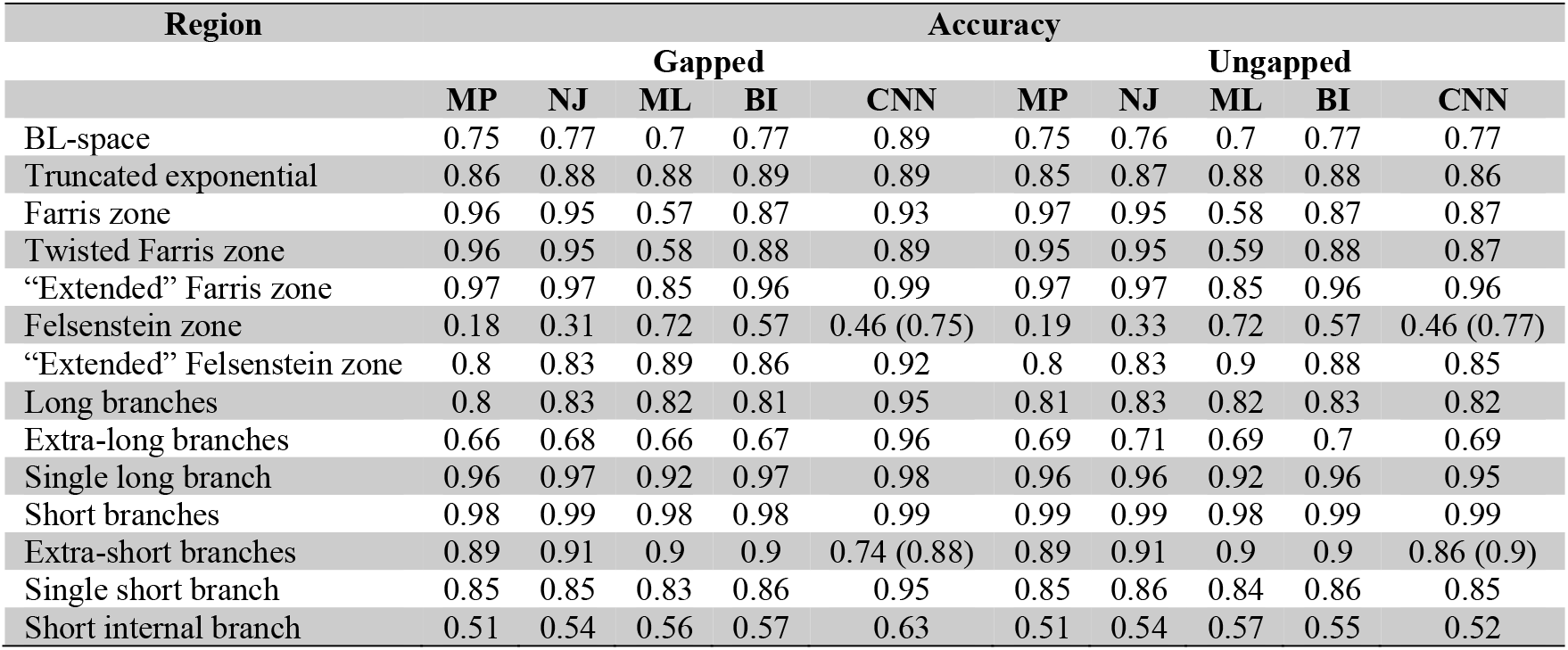
Performance accuracy of different topology estimation methods on simulated MSAs of length 1000. Accuracy estimates obtained from 15000 (5000 per unrooted topology) sampled from BL-space and from several additional test sets each consisting of 3000 MSAs (1000 per unrooted topology). The accuracy of CNN can be substantially improved by providing additional MSAs for training simulated from specific regions. We performed this additional training for the Felsenstein and extra-short branches regions and show the CNN’s improved accuracy in parentheses next to the original accuracy. MP=maximum parsimony, NJ=neighbor-joining, ML=maximum likelihood, BI=Bayesian inference, CNN=convolutional neuronal network.

Additionally, we tested performance of all methods on shorter gapped and ungapped MSAs of length 500 within each region specified in **Table 2** (see **Supplementary Table S1** for accuracy estimates). Overall, accuracies were lower for all methods, but with qualitatively similar rankings across methods. Further, we showed that accuracy increases with increasing MSA size for the CNN which is potentially indicative of statistical consistency (**Supplementary Table S2 and Supplementary Text**), though we do not prove this here.

Although we find that overall the CNN performs quite well on the test sets examined above, it is important to stress that it a method’s overall utility is not fully captured by a single accuracy estimate from one test set, or even a suite of test sets as we have done here. Each of the test sets we examined is a subspace of the BL-space that we defined in the **Supplementary Text**, and there are an infinite number of such sets that one could examine. Moreover, partitioning this space into a number of equally sized sets small enough to comprehensively assess how performance varies across the entire BL-space is a monumental task. For example, if one defined a BL-space and discretized it into 1% increments for all five branches, one would need to simulate 100^5^ datasets with all possible branch length combinations in order to traverse the entire parameter space. The unfeasibility of such an experiment was previously acknowledged by Huelsenbeck (1995), thus we cannot rule out the possibility that our CNN’s performance is compromised in certain regions of parameter space especially if examples from such regions were underrepresented during training phase. Indeed this appears be the cause of our initially poor performance on the Felsenstein and the “extra-short branches” zones, which was rectified through targeted additional training.

### Analysis of Phylogenetic Reliability Measures

Testing phylogenetic hypotheses usually involves calculation of some reliability measures such as bootstrap support (BS) for MP, NJ and ML or posterior probabilities (PP) in the context of BI. Despite the fact that their direct comparison is somewhat problematic due to their fundamental differences, one can ask which measures are more prone to support false tree topologies (Douady, et al. 2003; Huelsenbeck and Rannala 2004). For our CNN we used two confidence measures: bootstrap support for the chosen topology (obtained by applying the trained CNN to each MSA in a set of bootstrap replicates), and the class membership posterior probability (CPP) estimates produced automatically by the CNN which can also be interpreted as a measure of topological reliability. We find that high support (BS > 0.95, or PP > 0.95 in the case of BI) is occasionally assigned to incorrectly classified trees by several methods: 0.052 of misclassifications by MP, 0.05 by NJ, 0.003 by ML, 0.042 by BI, and 0.038 by the CNN for gapped MSAs (**Fig. 6a**) and 0.056 by MP, 0.054 by NJ, 0.0036 by ML, 0.048 by BI, and 0.034 by the CNN for ungapped MSAs (**Fig. 6b**). Thus, a very high reliability measure cutoff may be necessary in order to filter out positively misleading cases. On the other hand, the CPP estimates show a larger disparity between scores assigned to correctly vs. incorrectly classified topologies than do other measures (**Fig. 6**), and only 0.0004 and 0.0038 of incorrectly classified gapped and ungapped MSAs respectively having a CPP above 0.95. In other words, the CNN can avoid incorrect classifications even when imposing a relatively relaxed posterior probability cutoff, and thus simultaneously recover a large fraction of correctly classified topologies while producing very few misclassifications. This is further illustrated by precision-recall curves (**Supplementary Fig. S5**), which show that compared to other methods the CNN achieves high sensitivity while retaining a high positive predictive value on gapped alignments. Moreover, for our four-taxon topology trees the interpretation of CPP is straightforward: the most unreliable tree estimates receive CPP of ~1/3, which is equivalent to randomly picking one of the three possible topologies.

**Figure 6:**
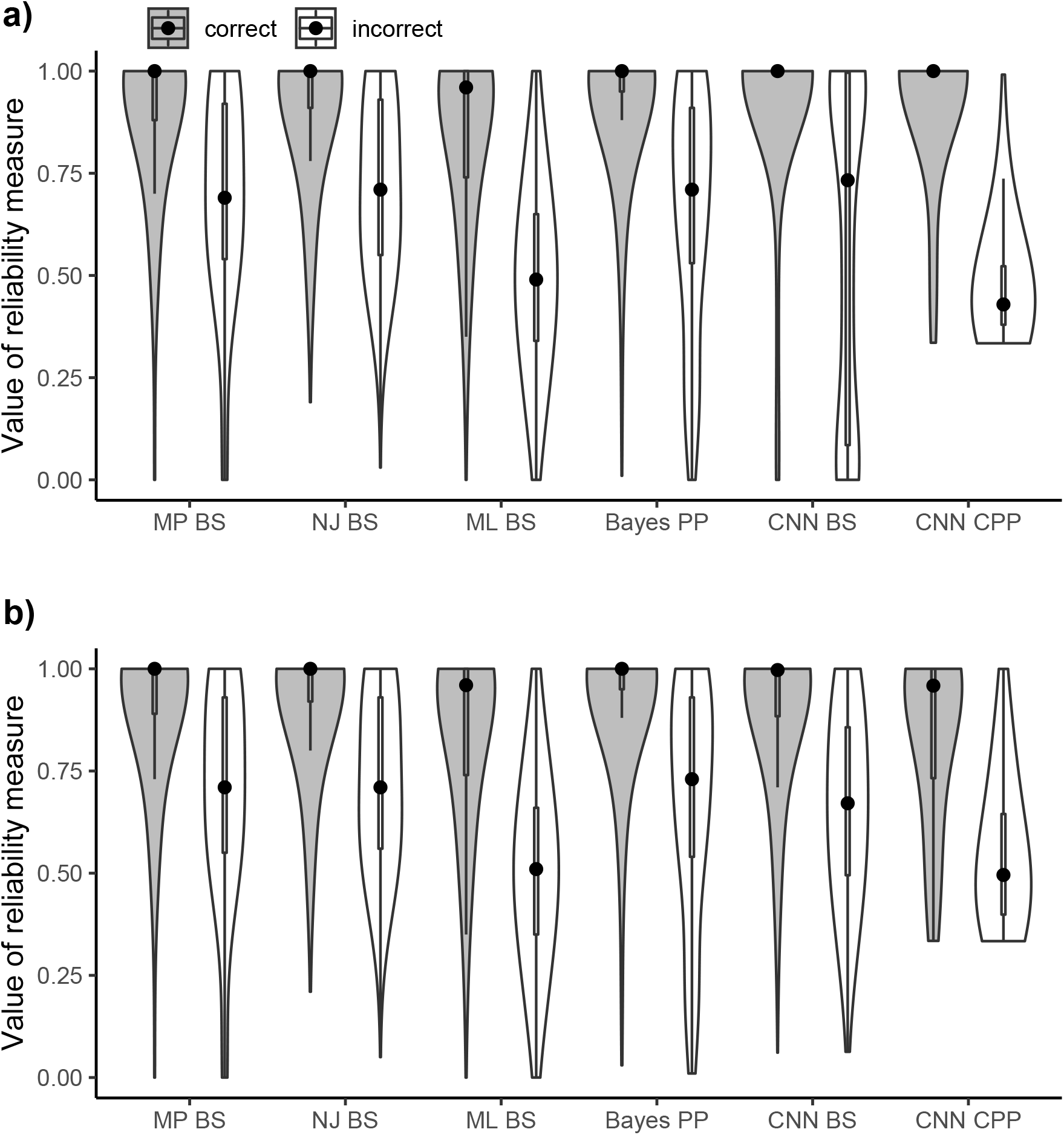
Comparisons of reliability measures among tree inference methods on gapped MSAs. The violin plots represent distribution of reliability measures (BS = bootstrap, PP = Bayesian posterior probability, CPP = CNN topology posterior probability) for correctly and incorrectly inferred topologies. MP=maximum parsimony, NJ=neighbor-joining, ML=maximum likelihood, BI=Bayesian inference, CNN=convolutional neuronal network.

### Concluding Remarks

We used a convolutional neural network to infer unrooted quartet topologies through classification. Our results demonstrate that CNNs perform this task quite well and have a number of desirable qualities. First and foremost, under many scenarios our CNN resolves quartets with higher accuracy than existing methods (**Table 2**). The CNN is also relatively insensitive to regions of the tree branch-length heterogeneity (e.g. the Felsenstein and Farris zones) that are known to confound certain methods, perhaps because the training set included trees with a variety of branch-length configurations (**see Supplementary Text**). Another advantage of using CNNs is that they produce posterior probability estimates for each tree topology; these posterior probabilities are estimated during the classification process and thus require no additional computation. Also, we have shown that our CNN’s accuracy can be further improved by adding more training datasets, albeit most likely with diminishing returns (**Fig. 2**). Although training on large datasets can impose a computational burden up front, this training only needs to be completed once, after which the CNN can be used to infer tree topologies very rapidly: the CNN can infer the topologies of 3000 MSAs of length 1000 in ~5 seconds using an NVIDIA Tesla V100-SXM2 GPU). Finally, CNNs have the advantage of being able to incorporate indels into their classification in a straightforward manner—in principle one need only include them in the simulated training data—which can improve accuracy by supplying more information to the classifier. However, future efforts will have to examine the robustness of inference to indel model misspecification, and the extent to which robustness can be improved by training under a mixture of indel rates and length distributions. More generally, this problem underscores the importance of considering ways in which empirical data may not match simulated data, and the consequences of such mismatches for inference. This problem is particularly challenging, as one may not always be able to predict the manner in which empirical data may differ substantially from model-based simulations, but could potentially be addressed by incorporating empirical data as well into the training process. For example, the CNN could be trained on simulated MSAs, and then a data set consisting of real MSAs for which there is strong consensus around the true topology can be supplied to the CNN for additional training iterations. Thus, through this transfer learning procedure (Pan and Yang 2010), the CNN would first learn to accurately infer trees from simulated MSA, and then be tuned to recognize some of the idiosyncrasies of real MSAs.

Our efforts thus suggest that there is great potential for the application of deep learning techniques to inferring trees, which is central to questions in both phylogenetics (species-tree inference) and population genomics (inferring ancestral recombination graphs along a recombining chromosome (Rasmussen, et al. 2014). Indeed, using the entire MSA as input would be most useful on recombining sequences—otherwise it is possible to collapse the MSA down to a vector of statistics (e.g. the number of occurrences of each distinct possible MSA column, if the number of sequences is small) that would sufficiently capture all of the relevant information from the input. Deep learning methods acting directly on the MSA could also be applied to additional phylogenetic problems, such as substitution model selection or tree inference from protein sequence data.

Although our findings are encouraging, additional advances are required for our approach to match the practical utility of conventional methods. First, we have only investigated the ability of a CNN to produce the correct tree topology, but not the associated branch lengths. Artificial neural networks are capable of estimating real-valued parameters in addition to discrete class labels, so future work should investigate the possibility of using deep learning for inferring branch lengths in addition to the topology. Another limitation of our current approach is that it is limited to a small number of sequences/taxa. Future extensions could use quartet amalgamation methods (e.g. von Haeseler and Strimmer 1996; Reaz, et al. 2014). Although such methods have not matched the accuracy of standard approaches such as ML in the past, because our method is able to resolve quartet topologies with greater accuracy and better calibrated confidence scores, it may be worth investigating whether quartet amalgamation combined via our CNN could compete with standard methods when applied to larger numbers of sequences. Alternatively, it may be possible to devise neural networks capable of directly predicting a tree topology. The latter could in principle be accomplished by treating the values of the final hidden layer of the network as entries in a distance matrix and using an output layer that builds a tree from this matrix (e.g. using NJ), and adopting a tree dissimilarity measure such as RF distance as the objective function to be minimized. However, this approach would also have a number of practical challenges that would have to be overcome (e.g. how to train networks that can handle different numbers of input sequences). One possible solution to this problem is to train a number of CNNs each designed to handle a fixed number of sequences, and then when given an input MSA select the CNN corresponding to the closest number of sequences greater than or equal to that of the input. The size of then input can then be increased by adding duplicates of one row of the MSA until the desired number of sequences is reached. After inferring the topology for this input, the duplicate nodes can be pruned from the tree to produce the final topology.

Finally, in order to move from inferring gene trees to inferring species trees, our approach will have to be extended to handle tree discordance due to variation in evolutionary rates across loci, incomplete lineage sorting (Maddison and Knowles 2006; Degnan and Rosenberg 2009) and gene flow (Maddison 1997). Supervised learning approaches may prove to be well suited for this question, as arbitrarily complicated phylogenetic models combining multiple sources of discordance can readily be incorporated into the training process via simulation. Thus, while additional practical developments are necessary, applications of deep learning to problems in tree inference and phylogenomics are likely to be a fruitful endeavor.

## Supplementary Material

Additional supplementary material is available at Dryad (DOI: XXX); R and Python scripts can be found at GitHub (https://github.com/SchriderLab/Tree_learning), also archived on Zenodo (DOI: 10.5281/zenodo.3246669). Training, validation and test MSA data sets are deposited on Figshare (https://doi.org/10.6084/m9.figshare.8279618.v2).

## Acknowledgements

We thank Matthew Hahn for valuable feedback on the manuscript, and Seth Bybee and Ariella Gladstein for helpful suggestions. We also thank Rob Lanfear and one anonymous reviewer for their comments.

## Literature Cited

Abadi M, Agarwal A, Barham P, Brevdo E, Chen Z, Citro C, Corrado GS, Davis A, Dean J, Devin M, et al. 2016. TensorFlow: Large-Scale Machine Learning on Heterogeneous Distributed Systems. eprint arXiv:1603.04467:arXiv:1603.04467.

Aberer AJ, Kobert K, Stamatakis A. 2014. ExaBayes: massively parallel bayesian tree inference for the whole-genome era. Molecular biology and evolution, 31:2553–2556.

Ashkenazy H, Cohen O, Pupko T, Huchon D. 2014. Indel reliability in indel-based phylogenetic inference. Genome biology and evolution, 6:3199–3209.

Bonham-Carter O, Steele J, Bastola D. 2013. Alignment-free genetic sequence comparisons: a review of recent approaches by word analysis. Briefings in bioinformatics.

Castresana J. 2000. Selection of conserved blocks from multiple alignments for their use in phylogenetic analysis. Molecular biology and evolution, 17:540–552.

Chan J, Perrone V, Spence JP, Jenkins PA, Mathieson S, Song YS. 2018. A Likelihood-Free Inference Framework for Population Genetic Data using Exchangeable Neural Networks. bioRxiv.

Degnan JH, Rosenberg NA. 2009. Gene tree discordance, phylogenetic inference and the multispecies coalescent. Trends in ecology & evolution, 24:332–340.

Dessimoz C, Gil M. 2010. Phylogenetic assessment of alignments reveals neglected tree signal in gaps. Genome biology, 11:R37.

Douady CJ, Delsuc F, Boucher Y, Doolittle WF, Douzery EJP. 2003. Comparison of Bayesian and Maximum Likelihood Bootstrap Measures of Phylogenetic Reliability. Molecular biology and evolution, 20:248–254.

Farris JS. 1970. Methods for Computing Wagner Trees. Syst Zool, 19:83–92.

Felsenstein J. 1978a. Cases in Which Parsimony or Compatibility Methods Will Be Positively Misleading. Syst Zool, 27:401–410.

Felsenstein J. 1978b. The Number of Evolutionary Trees. Syst Zool, 27:27–33.

Felsenstein J. 1981. Evolutionary trees from DNA sequences: a maximum likelihood approach. Journal of molecular evolution, 17:368–376.

Felsenstein J. 1985. Confidence-Limits on Phylogenies - an Approach Using the Bootstrap. Evolution; international journal of organic evolution, 39:783–791.

Felsenstein J. 1989. PHYLIP - Phylogeny Inference Package (Version 3.2). Cladistics, 5:164–166.

Fitch WM. 1971. Toward Defining the Course of Evolution: Minimum Change for a Specific Tree Topology. Syst Zool, 20:406–416.

Flagel L, Brandvain Y, Schrider DR. 2018. The Unreasonable Effectiveness of Convolutional Neural Networks in Population Genetic Inference. Molecular biology and evolution, 36:220–238.

Fletcher W, Yang Z. 2009. INDELible: a flexible simulator of biological sequence evolution. Molecular biology and evolution, 26:1879–1888.

Fujimoto MS, Suvorov A, Jensen NO, Clement MJ, Bybee SM. 2016. Detecting false positive sequence homology: a machine learning approach. BMC bioinformatics, 17:101.

Goodfellow I, Bengio Y, Courville A. 2016. Deep Learning. Adapt Comput Mach Le:1–775.

Gouy M, Guindon S, Gascuel O. 2010. SeaView version 4: A multiplatform graphical user interface for sequence alignment and phylogenetic tree building. Molecular biology and evolution, 27:221–224.

Hellmuth M, Wieseke N, Lechner M, Lenhof H-P, Middendorf M, Stadler PF. 2015. Phylogenomics with paralogs. Proceedings of the National Academy of Sciences, 112:2058.

Hohl M, Ragan MA. 2007. Is multiple-sequence alignment required for accurate inference of phylogeny? Syst Biol, 56:206–221.

Huelsenbeck JP. 1995. Performance of Phylogenetic Methods in Simulation. Syst Biol, 44:17–48.

Huelsenbeck JP, Hillis DM. 1993. Success of Phylogenetic Methods in the Four-Taxon Case. Syst Biol, 42:247–264.

Huelsenbeck JP, Rannala B. 2004. Frequentist Properties of Bayesian Posterior Probabilities of Phylogenetic Trees Under Simple and Complex Substitution Models. Syst Biol, 53:904–913.

Huelsenbeck JP, Ronquist F. 2001. MRBAYES: Bayesian inference of phylogenetic trees. Bioinformatics, 17:754–755.

Ioffe S, Szegedy C. 2015. Batch Normalization: Accelerating Deep Network Training by Reducing Internal Covariate Shift. eprint arXiv:1502.03167:arXiv:1502.03167.

Jukes TH, Cantor CR. 1969. CHAPTER 24 - Evolution of Protein Molecules. In: Munro HN editor. Mammalian Protein Metabolism, Academic Press, p. 21–132.

Kingma DP, Ba J. 2014. Adam: A Method for Stochastic Optimization. eprint arXiv:1412.6980:arXiv:1412.6980.

Kolaczkowski B, Thornton JW. 2009. Long-branch attraction bias and inconsistency in Bayesian phylogenetics. PloS one, 4:e7891.

Koonin EV. 2005. Orthologs, paralogs, and evolutionary genomics. Annual review of genetics, 39:309–338.

Krizhevsky A, Sutskever I, E. Hinton G. 2012. ImageNet Classification with Deep Convolutional Neural Networks.

Lecun Y, Bottou L, Bengio Y, Haffner P. 1998. Gradient-based learning applied to document recognition. Proceedings of the IEEE, 86:2278–2324.

Li S, Pearl DK, Doss H. 2000. Phylogenetic Tree Construction Using Markov Chain Monte Carlo. Journal of the American Statistical Association, 95:493–508.

Maddison WP. 1997. Gene Trees in Species Trees. Syst Biol, 46:523–536.

Maddison WP, Knowles LL. 2006. Inferring Phylogeny Despite Incomplete Lineage Sorting. Syst Biol, 55:21–30.

McTavish EJ, Steel M, Holder MT. 2015. Twisted trees and inconsistency of tree estimation when gaps are treated as missing data – The impact of model mis-specification in distance corrections. Molecular phylogenetics and evolution, 93:289–295.

Misof B, Misof K. 2009. A Monte Carlo approach successfully identifies randomness in multiple sequence alignments: a more objective means of data exclusion. Syst Biol, 58:21–34.

Nair V, E. Hinton G. 2010. Rectified Linear Units Improve Restricted Boltzmann Machines Vinod Nair.

Nguyen LT, Schmidt HA, von Haeseler A, Minh BQ. 2015. IQ-TREE: a fast and effective stochastic algorithm for estimating maximum-likelihood phylogenies. Molecular biology and evolution, 32:268–274.

Ogden TH, Rosenberg MS. 2007. How should gaps be treated in parsimony? A comparison of approaches using simulation. Molecular phylogenetics and evolution, 42:817–826.

Pan SJ, Yang Q. 2010. A Survey on Transfer Learning. IEEE Transactions on Knowledge and Data Engineering, 22:1345–1359.

Posada D. 2003. Using MODELTEST and PAUP* to Select a Model of Nucleotide Substitution. Current Protocols in Bioinformatics, 00:6.5.1–6.5.14.

Rannala B, Yang Z. 1996. Probability distribution of molecular evolutionary trees: a new method of phylogenetic inference. Journal of molecular evolution, 43:304–311.

Rasmussen MD, Hubisz MJ, Gronau I, Siepel A. 2014. Genome-Wide Inference of Ancestral Recombination Graphs. PLoS genetics, 10:e1004342.

Reaz R, Bayzid MS, Rahman MS. 2014. Accurate phylogenetic tree reconstruction from quartets: a heuristic approach. PloS one, 9:e104008–e104008.

Roch S. 2006. A short proof that phylogenetic tree reconstruction by maximum likelihood is hard. IEEE/ACM Transactions on Computational Biology and Bioinformatics, 3:92–94.

Rokas A, Holland PWH. 2000. Rare genomic changes as a tool for phylogenetics. Trends in ecology & evolution, 15:454–459.

Saitou N, Nei M. 1987. The neighbor-joining method: a new method for reconstructing phylogenetic trees. Molecular biology and evolution, 4:406–425.

Schrider DR, Kern AD. 2018. Supervised Machine Learning for Population Genetics: A New Paradigm. Trends in genetics: TIG, 34:301–312.

Siddall ME. 1998. Success of Parsimony in the Four-Taxon Case: Long-Branch Repulsion by Likelihood in the Farris Zone. Cladistics, 14:209–220.

Susko E. 2015. Bayesian long branch attraction bias and corrections. Syst Biol, 64:243–255.

Swofford DL, Waddell PJ, Huelsenbeck JP, Foster PG, Lewis PO, Rogers JS. 2001. Bias in Phylogenetic Estimation and Its Relevance to the Choice between Parsimony and Likelihood Methods. Syst Biol, 50:525–539.

Tarca AL, Carey VJ, Chen XW, Romero R, Draghici S. 2007. Machine learning and its applications to biology. PLoS computational biology, 3:e116.

Tavaré S. 1986. Some Probabilistic and Statistical Problems in the Analysis of DNA Sequences. American Mathematical Society: Lectures on Mathematics in the Life Sciences, Amer Mathematical Society, p. 57–86.

Truszkowski J, Goldman N. 2016. Maximum Likelihood Phylogenetic Inference is Consistent on Multiple Sequence Alignments, with or without Gaps. Syst Biol, 65:328–333.

Vialle RA, Tamuri AU, Goldman N. 2018. Alignment Modulates Ancestral Sequence Reconstruction Accuracy. Molecular biology and evolution, 35:1783–1797.

von Haeseler A, Strimmer K. 1996. Quartet Puzzling: A Quartet Maximum-Likelihood Method for Reconstructing Tree Topologies. Molecular biology and evolution, 13:964–964.

Warnow T. 2012. Standard maximum likelihood analyses of alignments with gaps can be statistically inconsistent. PLoS currents, 4:RRN1308.

Yang Z. 1994. Yang Z.. Estimating the pattern of nucleotide substitution. J Mol Evol 39: 105–111.

