## Supplementary figures and images for "Accurate inference of tree topologies from multiple sequence alignments using deep learning"

### Suppl Fig S1

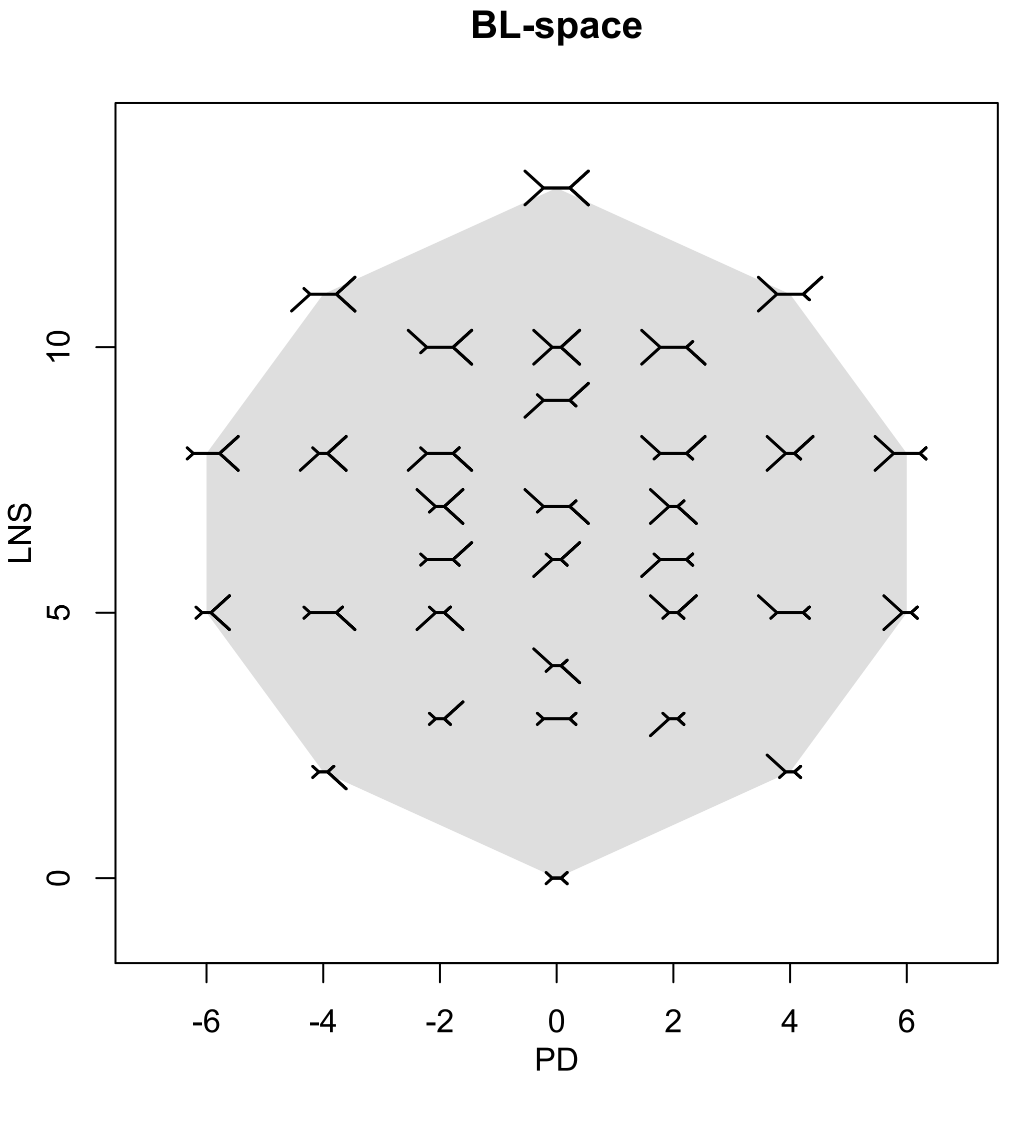

### Suppl Fig S2

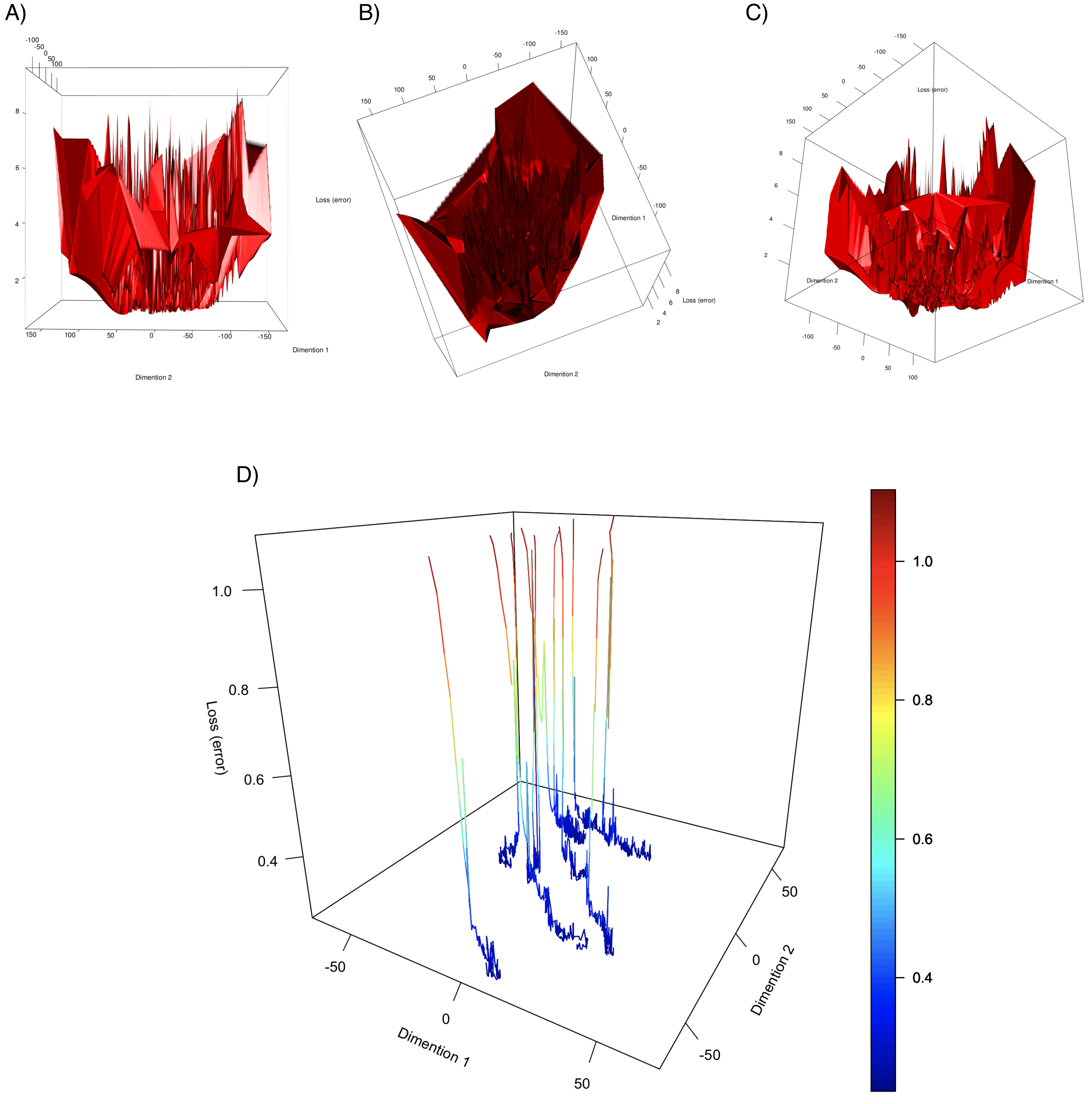

### Suppl Fig S3

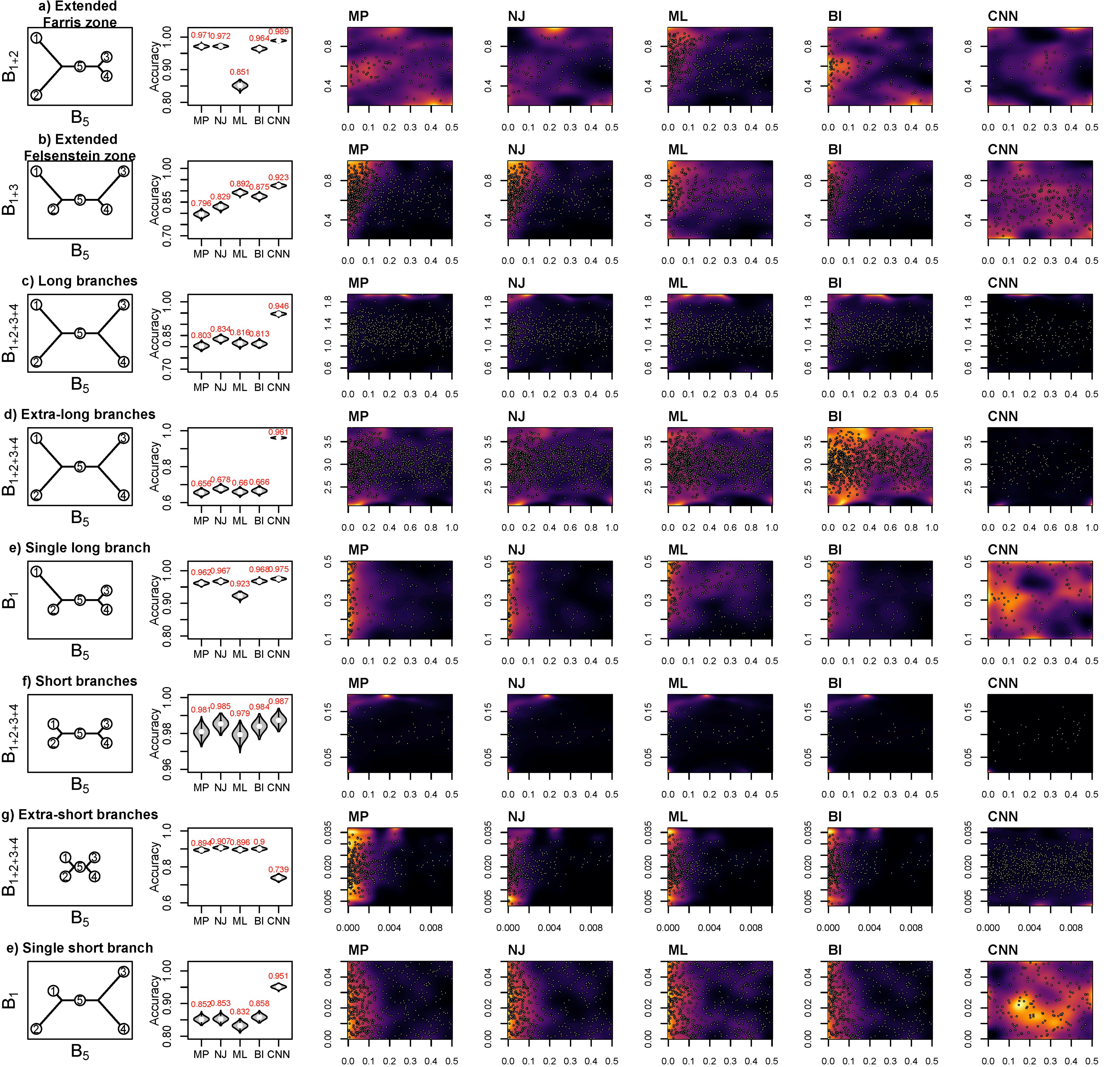

### Suppl Fig S4

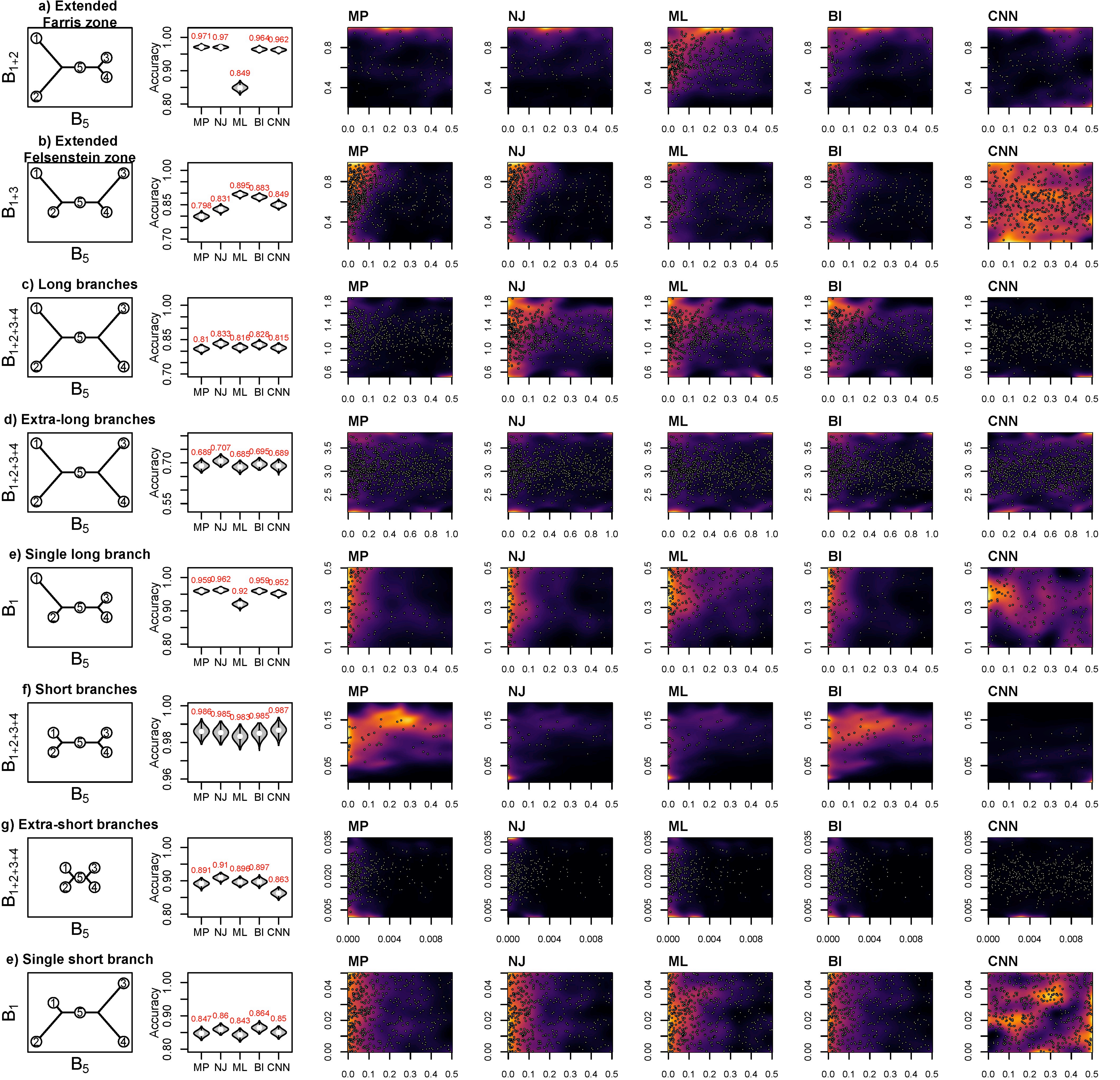

### Suppl Fig S5

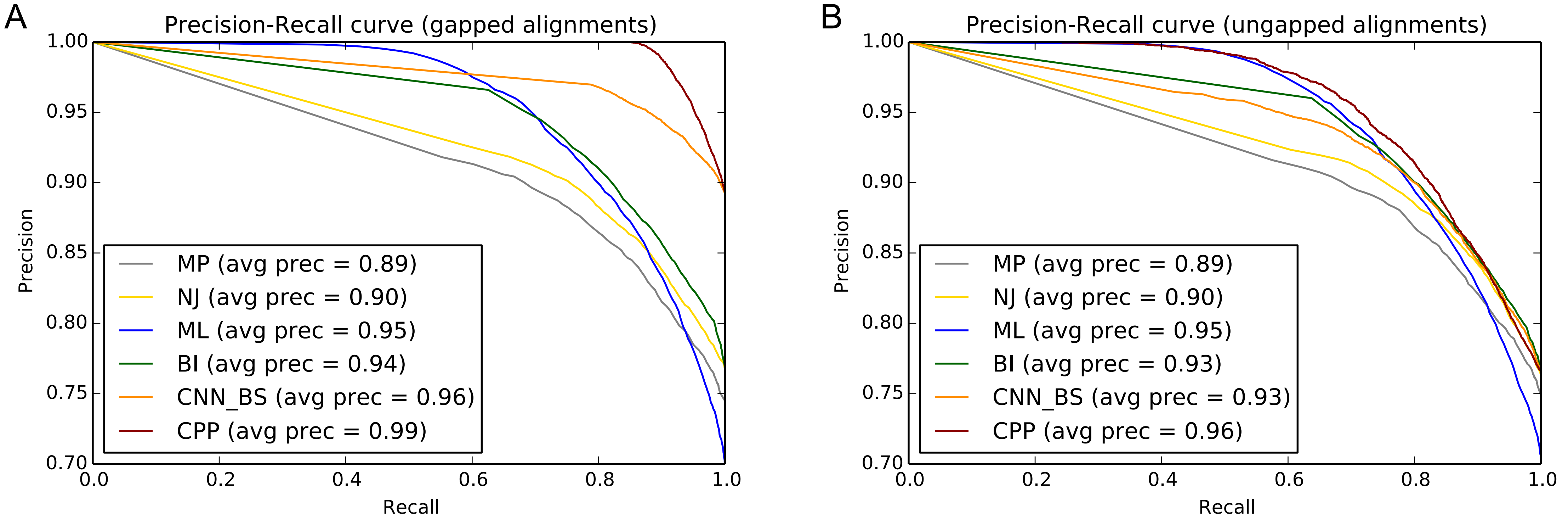
