## Supplementary material for "Accurate inference of tree topologies from multiple sequence alignments using deep learning": Suppl Tab S1

**Table S1: Performance accuracy of different topology estimation methods**. Accuracy estimates obtained from 15000 (5000 per unrooted topology) for BL-space and from 3000 (1000 per unrooted topology) for other regions simulated MSAs of length 500. The accuracy of CNN can be substantially improved by providing additional MSAs for training simulated from specific regions. MP=maximum parsimony, NJ=neighbor-joining, ML=maximum likelihood, BI=Bayesian inference, CNN=convolutional neuronal network.

| Region | **Accuracy** | | | | | | | | | |
| --- | --- | --- | --- | --- | --- | --- | --- | --- | --- | --- |
|  | **Gapped** | | | | | **Ungapped** | | | | |
|  | **MP** | **NJ** | **ML** | **BI** | **CNN** | **MP** | **NJ** | **ML** | **BI** | **CNN** |
| BL-space | 0.73 | 0.75 | 0.66 | 0.74 | 0.86 | 0.72 | 0.75 | 0.67 | 0.74 | 0.73 |
| Truncated exponential | 0.82 | 0.84 | 0.84 | 0.75 | 0.89 | 0.84 | 0.85 | 0.84 | 0.83 | 0.83 |
| Farris zone | 0.94 | 0.92 | 0.53 | 0.86 | 0.77 | 0.94 | 0.92 | 0.53 | 0.85 | 0.87 |
| Twisted Farris zone | 0.92 | 0.91 | 0.55 | 0.84 | 0.74 | 0.94 | 0.91 | 0.56 | 0.84 | 0.82 |
| “Extended” Farris zone | 0.96 | 0.96 | 0.82 | 0.95 | 0.96 | 0.96 | 0.96 | 0.81 | 0.95 | 0.96 |
| Felsenstein zone | 0.21 | 0.31 | 0.65 | 0.48 | 0.41 | 0.2 | 0.31 | 0.66 | 0.48 | 0.35 |
| “Extended” Felsenstein zone | 0.78 | 0.82 | 0.87 | 0.86 | 0.88 | 0.78 | 0.82 | 0.88 | 0.86 | 0.81 |
| Long branches | 0.77 | 0.79 | 0.77 | 0.78 | 0.89 | 0.77 | 0.79 | 0.78 | 0.78 | 0.77 |
| Extra-long branches | 0.6 | 0.62 | 0.6 | 0.61 | 0.9 | 0.61 | 0.66 | 0.63 | 0.64 | 0.63 |
| Single long branch | 0.95 | 0.96 | 0.91 | 0.95 | 0.94 | 0.94 | 0.95 | 0.9 | 0.95 | 0.94 |
| Short branches | 0.97 | 0.98 | 0.94 | 0.98 | 0.94 | 0.97 | 0.97 | 0.93 | 0.97 | 0.97 |
| Extra-short branches | 0.83 | 0.83 | 0.82 | 0.83 | 0.38 | 0.82 | 0.83 | 0.83 | 0.83 | 0.79 |
| Single short branch | 0.81 | 0.84 | 0.81 | 0.83 | 0.9 | 0.82 | 0.83 | 0.8 | 0.83 | 0.83 |
| Short internal branch | 0.48 | 0.52 | 0.52 | 0.51 | 0.51 | 0.46 | 0.51 | 0.5 | 0.55 | 0.48 |
