## Supplementary material for "Accurate inference of tree topologies from multiple sequence alignments using deep learning": Suppl Tab S2

**Table S2: Accuracy estimates for different alignment lengths within BL-space**. Accuracy estimates obtained from 15000 (5000 per unrooted topology) simulated MSAs of length 500, 1000 or 10000 for different methods. MP=maximum parsimony, NJ=neighbor-joining, ML=maximum likelihood, BI=Bayesian inference, CNN=convolutional neuronal network.

| Method | **Accuracy** | | | | | |
| --- | --- | --- | --- | --- | --- | --- |
|  | **Gapped** | | | **Ungapped** | | |
|  | **500** | **1000** | **10000** | **500** | **1000** | **10000** |
| MP | 0.73 | 0.75 | 0.81 | 0.72 | 0.75 | 0.81 |
| NJ | 0.75 | 0.77 | 0.81 | 0.75 | 0.76 | 0.82 |
| ML | 0.66 | 0.7 | 0.79 | 0.67 | 0.7 | 0.80 |
| BI | 0.74 | 0.77 | 0.84 | 0.74 | 0.77 | 0.84 |
| CNN | 0.86 | 0.89 | 0.92 | 0.73 | 0.77 | 0.92 |
