## Supplementary Text for "Accurate inference of tree topologies from multiple sequence alignments using deep learning"

*Simulation from "the Branch-Length Space" (BL-space)*

The goal of our simulation strategy for constructing our training set was to generate data set consisting of a wide variety of tree branch-length configurations, each given equal representation. Simply drawing each branch length uniformly is not sufficient for this, as some tree configurations will be underrepresented (e.g. those where *all* five branches are short or *all* five branches are long) relative to others (e.g. those with a mixture of short, medium, and long branches). Thus we first generated a large number of simulations encompassing many different possible branch-length configurations (but not necessarily uniformly), projected them onto a branch-length space that we created to give more equal representation to these configurations (**Supplementary Fig. S1**), before drawing uniformly across this space. We describe each of these steps in detail below.

To begin, we generated a large number of tree topologies by randomly drawing the length (expressed in expected number of substitutions per site) of each branch (B) from the following mixture of beta distributions:

$$b \sim w_1 Beta(\alpha = \beta = 0.1) + w_2 Beta(\alpha = \beta = 0.5) + w_3 Beta(\alpha = \beta = 1),$$
$$w_1 = w_2 = w_3 = 1/3$$

Note that branch lengths generated from the beta components with $\alpha = \beta < 1$ produce extreme tree branch lengths (i.e. close to 0 or 1) with higher probability, whereas the beta component with $\alpha = \beta = 1$ is equivalent to standard uniform distribution. Thus, the total branch length of a tree ranged from 0 to 4 substitutions per site.

Although this strategy is useful for generating a wide variety of tree branch-length configurations, it does not generate all possible combinations of branch lengths in a uniform manner. Thus, we sought to subsample from this distribution in order to sample tree branch-length configurations more uniformly. To this end, for each simulated tree we projected the vector of branch lengths from $\mathbb{R}^5$ onto a $\mathbb{R}^2$ coordinate space, i.e. projecting 5D space where each dimension corresponds to a tree branch onto 2D space. To achieve this we used three simple statistics: the sum of pairwise differences (PD) of branch lengths, total tree length (L) and sum of lengths of neighboring branches (NS). These statistics were defined as follows:

$$\mathrm{PD} = \sum_{i<j} (B_i - B_j)$$
$$\mathrm{L} = \sum_{i} B_i$$
$$\mathrm{NS} = \sum_{i<5} (B_i + B_{i+1})$$

where $B_i$ denotes the length of the $i^{\text{th}}$ branch. The PD and L+NS (LNS) scores were used as *x* and *y* coordinates respectively for mapping on $\mathbb{R}^2$ that we termed "the branch length space" (BL-space). Although these statistics are somewhat arbitrary in their construction, we found they effectively separated the different possible tree branch-length configurations when visualized in two-dimensional space. For example, if we consider all branch-length configurations where each branch is either very short (i.e. close to zero) or very large (close to 1), there are $2^5$=32 possible arrangements, all of which are visible in this space (**Supplementary Fig. S1)**. On the other hand, alternative branch-length spaces that we considered, such as the branch-length space proposed in (Huelsenbeck and Hillis 1993) where tree nodal rotation is not considered, caused some of these 32 arrangements to be collapsed onto one another and thus could not represent as diverse a set of branch-length configurations.

In total, we simulated $10^8$ trees with different branch-length configurations using the aforementioned beta mixture to cover our entire BL-space and then sampled uniformly across this space. This was done by partitioning the BL-space shown in **Supplementary Fig. S1** into a grid of cells (each sized 0.1 × 0.1) and sampling an equal number of trees from each of these cells. This sampling scheme thus yields a wide variety of tree branch-length configurations, each equally represented, in our training data.

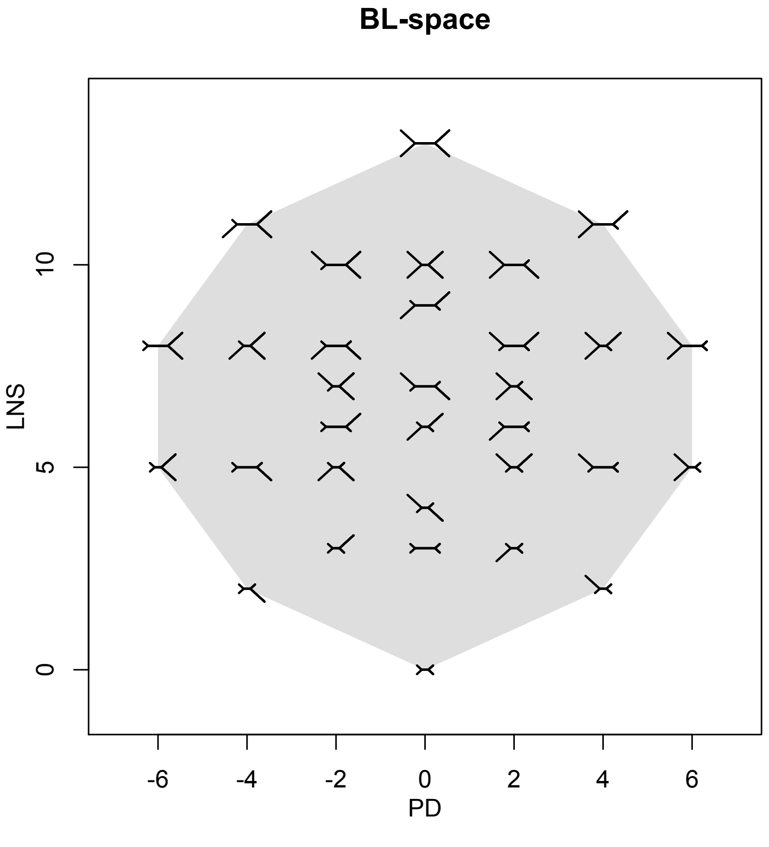


**Figure S1: The analytical branch length space (BL-space).** Each tree in the BL-space marks the location of the extreme cases where short branches in the image represent branch lengths equal to 0 and long branches represent lengths of 1. The shaded area shows the boundaries of the BL-space. PD = pairwise difference, LNS = tree length (L) + lengths of neighboring branches (NS).

*Visualizing the error surface during CNN training*

In order to visualize the error (or loss) surface of our CNN (**Supplementary Fig. S2**), we sampled different parameter combinations (i.e. neural network weights) in three different ways and evaluated the loss function of the CNN for each set of weights. We then reduced these sets of parameters to just two dimensions (see below) and plotted these two dimensions against their associated error value to produce a 3D plot of the error surface of our network. Our different sampling approaches capture different and somewhat complementary information, so our final plots pooled points sampled from each of three methods together. The code used compute points and to produce the plots is available online at https://github.com/SchriderLab/error_surface.

For the first sampling method, parameter values were drawn from (-0.05, 0.05). This is analogous to a random weight initialization step for the CNN and provides information about the possible starting points for training. The second sampling method is an adaptation of the method proposed by (Goodfellow, et al. 2014), which was used to visualize the error along the path from the initial to final trained value of the network. We began by training the network once to produce a low-loss set of parameters, and then generated a single point drawn from $U(-0.05, 0.05)$. New points were sampled along a straight line between the random point and the final point after training, with increasing density closer to the trained point. The same process was then performed on a line from the trained point moving in the direction opposite the random point. This resulted in a line bisected by the trained point, with random weight combinations and their associated values of the loss function sampled in the vicinity of this line, and with especially dense sampling near its center. This whole process was repeated 100 times for the same trained point, resulting in 100 different lines with 40 samples along each line. This sampling method is intended to provide a local context for a trained point, and characterize the parts of the loss surface that must be navigated by the optimization algorithm, with higher resolution in the vicinity of final trained parameterization (which is often a local optimum). For the final sampling method, we recorded the parameters for a network at regular intervals during training, thereby showing the trajectory in weight space for a particular training run.

The CNN architectures that we explored have on the order of one million weights, so reducing the dimensionality of this weight space to just two is not a trivial transformation. Principal components analysis (PCA) is not optimal for our problem due to performance issues. Conventional implementations PCA require that all samples be in memory at the time of calculation, which severely limits the number of sampled sets of weights that we can produce for a given network. As an alternative, we implemented sparse random matrix projections (Li, et al. 2006). Sparse random matrix projections have a number of advantages over PCA for our purposes. First, the transformation matrix can be generated before sampling, because only knowledge of the starting dimensions and desired final dimensions is required. Second, the transformations can be calculated in a streaming fashion, with each sampled point reduced to two dimensions before being saved to disk. This requires only one set of parameters to be in memory at a time. Third, the transformation to a lower dimensional embedding can be preserved after the completion of the process by saving the transformation matrix. This allows flexibility in sampling, which can be performed in different processes or even on different machines. We found that the computational bottleneck for our visualization process was the loss evaluation for a given set of weights, which depends on GPU-accelerated calculations. The flexibility of sparse random matrix projection allows the use of GPUs across different machines without needing shared memory. It is important to note that there is still a significant amount of information lost in this transformation process, but this is an unavoidable problem associated with reducing such a high-dimensional space to just two dimensions.

*Classification Accuracy Increases with Alignment Length*

Statistical consistency is defined as the ability of a method, $M$, to return the true parameter value, $\theta$ (e.g. tree topology), with probability approaching unity when amount of data ($D$) increases (with alignment length measured in nucleotides, $n$). More formally, a method is consistent when:

$$\lim_{n\to\infty} P(M(D_n) = \theta) = 1$$

Statistical consistency has been considered as one of the major criteria for a good tree reconstruction method (Felsenstein 1978; Goldman 1990; Penny, et al. 1992). Consistency of machine learning models has been only formally proven for a multilayer perceptron regression estimator with a particular network architecture (Mielniczuk and Tyrcha 1993). Generally, it is not trivial to analytically derive the global theoretical guarantees for neural networks due to their complexity and flexibility as exemplified by their large number of hyperparameters (e.g. different network architectures, activation and loss functions and optimization algorithms), which can be tuned to accommodate a certain learning problem. Additional factors such as sizes of both the training dataset and the input data (in our case number of sequences in the MSA and the MSA's length) may also affect the degree of consistency. Here we considered three consistent methods (provided the correct substitution model is used) of conventional tree estimation: NJ (Saitou and Nei 1987), ML (Felsenstein 1981; Truszkowski and Goldman 2016) and BI (Steel, et al. 2013; Warnow 2018) as well as MP as an example of an inconsistent (Felsenstein 1978) method under standard Markov substitution models. Because empirical datasets are always finite in length, the rate (Fu 1975) at which $P(M(D_n) = \theta)$ increases with $n$ is perhaps more crucial in practice.

We tested our CNN and the four standard methods on simulated sets of MSAs of increasing length. Our results show (**Supplementary Table S2**) that for both gapped and ungapped MSAs each method's accuracy increases with MSA length, and that the CNN outperforms other methods for MSA of length 10000. Thus, although we do not prove that CNNs are consistent, these empirical results are highly encouraging and imply that our approach has great potential for practical applications. Indeed, in some instances accuracy on shorter MSAs may be a more crucial quality than provable consistency because asymptotic properties of consistent methods may not be exhibited for smaller datasets (Yang 2014).

*Tree inference commands*

<u>Maximum parsimony (MP) using PHYLIP implemented in seaview</u>:

```
seaview -build_tree -parsimony -o - -unroot -gaps_as_unknown -
search more -replicates 1000 file.fasta
```

<u>Maximum likelihood (ML) using IQTREE</u>:

1) Model selection

```
iqtree -s file.fasta -mset JC,TIMe,TIM,GTR,12.12 -mrate
G8,I,I+G8 -blmin 1e-100 -b 1000 -quiet -pre ML -safe
```

2) Exhaustive ML search

```
iqtree -s file.fasta -te tree1 -m best_model -blmin 1e-100 -quiet -pre tree1 -safe
iqtree -s file.fasta -te tree2 -m best_model -blmin 1e-100 -quiet -pre tree2 -safe
iqtree -s file.fasta -te tree3 -m best_model -blmin 1e-100 -quiet -pre tree3 -safe
```

`tree1`, `tree2` and `tree3` are the text files that contain ((A,D),B,C), ((B,D),A,C) and ((C,D),A,B) topologies in newick format, respectively.

Neighbor joining (NJ) using IQTREE:

```
iqtree -s file.fasta -b 1000 -t BIONJ -m best_model -ninit 0 -ntop 1 -n 0 -blfix -pre NJ -safe
```

Bayesian inference (BI) using ExaBayes:

```
yggdrasil -f fasta.phy -m DNA -n myRun -s 77 -c config.nex
consense -f ExaBayes_topologies.myRun.* -n myCons
```

Contents of the config.nex file:

```
#NEXUS

begin run;
   parsimonyStart false
   proposalSets true
   numCoupledChains 2
   numGen 1e6
   numRuns 3
end;
```

*Main CNN network architectures (Keras)*

1) Gapped alignment; 50k MSAs per topology; MSA length 1000 (note indels make MSA larger)

```
Train on 150000 samples, validate on 15000 samples

Layer (type)                 Output Shape              Param #
=================================================================
input_1 (InputLayer)         (None, 4, 1596, 1)        0
_________________________________________________________________
zero_padding2d_1 (ZeroPaddin (None, 4, 1596, 1)        0
_________________________________________________________________
conv2d_1 (Conv2D)            (None, 1, 1596, 1024)     5120
_________________________________________________________________
batch_normalization_1 (Batch (None, 1, 1596, 1024)     4096
```

| | | |
|---|---|---|
| dropout_1 (Dropout) | (None, 1, 1596, 1024) | 0 |
| average_pooling2d_1 (Average | (None, 1, 1596, 1024) | 0 |
| zero_padding2d_2 (ZeroPaddin | (None, 1, 1597, 1024) | 0 |
| conv2d_2 (Conv2D) | (None, 1, 1596, 1024) | 2098176 |
| batch_normalization_2 (Batch | (None, 1, 1596, 1024) | 4096 |
| dropout_2 (Dropout) | (None, 1, 1596, 1024) | 0 |
| average_pooling2d_2 (Average | (None, 1, 399, 1024) | 0 |
| zero_padding2d_3 (ZeroPaddin | (None, 1, 400, 1024) | 0 |
| conv2d_3 (Conv2D) | (None, 1, 399, 128) | 262272 |
| batch_normalization_3 (Batch | (None, 1, 399, 128) | 512 |
| dropout_3 (Dropout) | (None, 1, 399, 128) | 0 |
| average_pooling2d_3 (Average | (None, 1, 99, 128) | 0 |
| zero_padding2d_4 (ZeroPaddin | (None, 1, 100, 128) | 0 |
| conv2d_4 (Conv2D) | (None, 1, 99, 128) | 32896 |
| batch_normalization_4 (Batch | (None, 1, 99, 128) | 512 |
| dropout_4 (Dropout) | (None, 1, 99, 128) | 0 |
| average_pooling2d_4 (Average | (None, 1, 24, 128) | 0 |
| zero_padding2d_5 (ZeroPaddin | (None, 1, 25, 128) | 0 |
| conv2d_5 (Conv2D) | (None, 1, 24, 128) | 32896 |
| batch_normalization_5 (Batch | (None, 1, 24, 128) | 512 |
| dropout_5 (Dropout) | (None, 1, 24, 128) | 0 |
| average_pooling2d_5 (Average | (None, 1, 12, 128) | 0 |
| zero_padding2d_6 (ZeroPaddin | (None, 1, 13, 128) | 0 |
| conv2d_6 (Conv2D) | (None, 1, 12, 128) | 32896 |
| batch_normalization_6 (Batch | (None, 1, 12, 128) | 512 |
| dropout_6 (Dropout) | (None, 1, 12, 128) | 0 |
| average_pooling2d_6 (Average | (None, 1, 6, 128) | 0 |
| zero_padding2d_7 (ZeroPaddin | (None, 1, 7, 128) | 0 |

```
conv2d_7 (Conv2D)            (None, 1, 6, 128)         32896     
_________________________________________________________________
batch_normalization_7 (Batch (None, 1, 6, 128)         512       
_________________________________________________________________
dropout_7 (Dropout)          (None, 1, 6, 128)         0         
_________________________________________________________________
average_pooling2d_7 (Average (None, 1, 3, 128)         0         
_________________________________________________________________
zero_padding2d_8 (ZeroPaddin (None, 1, 4, 128)         0         
_________________________________________________________________
conv2d_8 (Conv2D)            (None, 1, 3, 128)         32896     
_________________________________________________________________
batch_normalization_8 (Batch (None, 1, 3, 128)         512       
_________________________________________________________________
dropout_8 (Dropout)          (None, 1, 3, 128)         0         
_________________________________________________________________
average_pooling2d_8 (Average (None, 1, 3, 128)         0         
_________________________________________________________________
flatten_1 (Flatten)          (None, 384)               0         
_________________________________________________________________
dense_1 (Dense)              (None, 1024)              394240    
_________________________________________________________________
dropout_9 (Dropout)          (None, 1024)              0         
_________________________________________________________________
dense_2 (Dense)              (None, 3)                 3075      
=================================================================
Total params: 2,938,627
Trainable params: 2,932,995
Non-trainable params: 5,632
_________________________________________________________________
```

### 2) Ungapped alignment; 50k MSAs per topology; MSA length 1000

```
Train on 150000 samples, validate on 15000 samples

Layer (type)                 Output Shape              Param #   
=================================================================
input_1 (InputLayer)         (None, 4, 1000, 1)        0         
_________________________________________________________________
zero_padding2d_1 (ZeroPaddin (None, 4, 1000, 1)        0         
_________________________________________________________________
conv2d_1 (Conv2D)            (None, 1, 1000, 1024)     5120      
_________________________________________________________________
batch_normalization_1 (Batch (None, 1, 1000, 1024)     4096      
_________________________________________________________________
dropout_1 (Dropout)          (None, 1, 1000, 1024)     0         
_________________________________________________________________
average_pooling2d_1 (Average (None, 1, 1000, 1024)     0         
_________________________________________________________________
zero_padding2d_2 (ZeroPaddin (None, 1, 1001, 1024)     0         
_________________________________________________________________
conv2d_2 (Conv2D)            (None, 1, 1000, 1024)     2098176   
_________________________________________________________________
batch_normalization_2 (Batch (None, 1, 1000, 1024)     4096      
_________________________________________________________________
```

```
dropout_2 (Dropout)          (None, 1, 1000, 1024)     0
_________________________________________________________________
average_pooling2d_2 (Average (None, 1, 250, 1024)      0
_________________________________________________________________
zero_padding2d_3 (ZeroPaddin (None, 1, 251, 1024)      0
_________________________________________________________________
conv2d_3 (Conv2D)            (None, 1, 250, 128)       262272
_________________________________________________________________
batch_normalization_3 (Batch (None, 1, 250, 128)       512
_________________________________________________________________
dropout_3 (Dropout)          (None, 1, 250, 128)       0
_________________________________________________________________
average_pooling2d_3 (Average (None, 1, 62, 128)        0
_________________________________________________________________
zero_padding2d_4 (ZeroPaddin (None, 1, 63, 128)        0
_________________________________________________________________
conv2d_4 (Conv2D)            (None, 1, 62, 128)        32896
_________________________________________________________________
batch_normalization_4 (Batch (None, 1, 62, 128)        512
_________________________________________________________________
dropout_4 (Dropout)          (None, 1, 62, 128)        0
_________________________________________________________________
average_pooling2d_4 (Average (None, 1, 15, 128)        0
_________________________________________________________________
zero_padding2d_5 (ZeroPaddin (None, 1, 16, 128)        0
_________________________________________________________________
conv2d_5 (Conv2D)            (None, 1, 15, 128)        32896
_________________________________________________________________
batch_normalization_5 (Batch (None, 1, 15, 128)        512
_________________________________________________________________
dropout_5 (Dropout)          (None, 1, 15, 128)        0
_________________________________________________________________
average_pooling2d_5 (Average (None, 1, 7, 128)         0
_________________________________________________________________
zero_padding2d_6 (ZeroPaddin (None, 1, 8, 128)         0
_________________________________________________________________
conv2d_6 (Conv2D)            (None, 1, 7, 128)         32896
_________________________________________________________________
batch_normalization_6 (Batch (None, 1, 7, 128)         512
_________________________________________________________________
dropout_6 (Dropout)          (None, 1, 7, 128)         0
_________________________________________________________________
average_pooling2d_6 (Average (None, 1, 3, 128)         0
_________________________________________________________________
zero_padding2d_7 (ZeroPaddin (None, 1, 4, 128)         0
_________________________________________________________________
conv2d_7 (Conv2D)            (None, 1, 3, 128)         32896
_________________________________________________________________
batch_normalization_7 (Batch (None, 1, 3, 128)         512
_________________________________________________________________
dropout_7 (Dropout)          (None, 1, 3, 128)         0
_________________________________________________________________
average_pooling2d_7 (Average (None, 1, 1, 128)         0
_________________________________________________________________
zero_padding2d_8 (ZeroPaddin (None, 1, 2, 128)         0
_________________________________________________________________
conv2d_8 (Conv2D)            (None, 1, 1, 128)         32896
```

```
_________________________________________________________________
batch_normalization_8 (Batch (None, 1, 1, 128)         512
_________________________________________________________________
dropout_8 (Dropout)          (None, 1, 1, 128)         0
_________________________________________________________________
average_pooling2d_8 (Average (None, 1, 1, 128)         0
_________________________________________________________________
flatten_1 (Flatten)          (None, 128)               0
_________________________________________________________________
dense_1 (Dense)              (None, 1024)              132096
_________________________________________________________________
dropout_9 (Dropout)          (None, 1024)              0
_________________________________________________________________
dense_2 (Dense)              (None, 3)                 3075
=================================================================
Total params: 2,676,483
Trainable params: 2,670,851
Non-trainable params: 5,632
_________________________________________________________________
```

*CNN training runtimes*

On NVIDIA Tesla V100-SXM2 GPU
Gap 50k 500 bp: ~1.25h
Gap 50k 1000 bp: ~4.5h

**Literature Cited**


Felsenstein J. 1978. Cases in Which Parsimony or Compatibility Methods Will Be Positively Misleading. Syst Zool, 27:401-410.

Felsenstein J. 1981. Evolutionary trees from DNA sequences: a maximum likelihood approach. Journal of molecular evolution, 17:368-376.

Fu J. 1975. The Rate of Convergence of Consistent Point Estimators.

Goldman N. 1990. Maximum-Likelihood Inference of Phylogenetic Trees, with Special Reference to a Poisson-Process Model of DNA Substitution and to Parsimony Analyses. Syst Zool, 39:345-361.

Goodfellow IJ, Vinyals O, Saxe AM. 2014. Qualitatively characterizing neural network optimization problems. eprint arXiv:1412.6544:arXiv:1412.6544.

Huelsenbeck JP, Hillis DM. 1993. Success of Phylogenetic Methods in the Four-Taxon Case. Syst Biol, 42:247-264.

Li P, Hastie TJ, Church KW. 2006. Very sparse random projections. Proceedings of the 12th ACM SIGKDD international conference on Knowledge discovery and data mining. Philadelphia, PA, USA, ACM, p. 287-296.

Mielniczuk J, Tyrcha J. 1993. Consistency of Multilayer Perceptron Regression-Estimators. Neural Networks, 6:1019-1022.

Penny D, Hendy MD, Steel MA. 1992. Progress with methods for constructing evolutionary trees. Trends in ecology & evolution, 7:73-79.
Saitou N, Nei M. 1987. The neighbor-joining method: a new method for reconstructing phylogenetic trees. Molecular biology and evolution, 4:406-425.
Steel M, Linz S, Huson DH, Sanderson MJ. 2013. Identifying a species tree subject to random lateral gene transfer. Journal of theoretical biology, 322:81-93.
Truszkowski J, Goldman N. 2016. Maximum Likelihood Phylogenetic Inference is Consistent on Multiple Sequence Alignments, with or without Gaps. Syst Biol, 65:328-333.
Warnow T. 2018. Computational phylogenetics : an introduction to designing methods for phylogeny estimation. Cambridge University Press.
Yang Z. 2014. Molecular Evolution: A Statistical Approach. Molecular Evolution: A Statistical Approach:1-492.
